# A network medicine framework for multi-modal data integration in therapeutic target discovery

**DOI:** 10.1101/2025.05.29.656813

**Authors:** Greta Baltušytė, Isaac Toleman, James O. Jones, Sarah J. Welsh, Grant D. Stewart, Thomas J. Mitchell, Kourosh Saeb-Parsy, Namshik Han

## Abstract

The high cost and attrition rate of drug development underscore the need for more effective strategies for therapeutic target discovery. Here, we present a network medicine-based machine learning framework that integrates single-cell transcriptomics, bulk multi-omic profiles, genome-wide CRISPR perturbation screens, and protein-protein interaction networks to systematically prioritise disease-specific targets. Applied to clear cell renal cell carcinoma, the framework successfully recovered established targets and predicted five therapeutic candidates, with subsequent in vitro validation demonstrating that among these, ENO2 inhibition had the strongest anti-tumour effect, followed by LRRK2, a repurposing candidate with phase III Parkinson’s disease inhibitors. The proposed approach advances target discovery by moving beyond single-feature, single-modality heuristics to a scalable, machine learning-driven strategy that is generalisable across diseases.

## Introduction

Targeted therapies have reshaped the clinical management of cancer by enabling selective inhibition of disease-driving molecular alterations, offering improved efficacy and reduced systemic toxicity compared to conventional chemotherapeutics. Yet, despite their transformative potential, the promise of precision oncology remains constrained. A limited pool of actionable targets, tumour heterogeneity and the frequent development of drug resistance continue to restrict the long-term success of targeted interventions, underscoring the need for therapeutic landscape expansion.

Conventional approaches to target discovery have typically relied on single-gene hypotheses, often informed by mutational profiling or differential expression analyses. While these methods have uncovered key oncogenic drivers, they fall short in capturing the interplay of molecular interactions that underpin disease pathophysiology. Mounting evidence suggests that pathologies such as cancer emerge not from isolated molecular events but from the disruption of coordinated gene networks [1–3]. This systems-level perspective has led to the emergence of network medicine, which leverages insights from network theory [4–6] to model the molecular architecture of disease. Among the most important observations in this field is that disease-associated genes often cluster in the same network neighbourhood, forming a disease module. Empirical studies have confirmed that drug targets tend to reside within or near these modules [7, 8], suggesting that network topology can be harnessed to prioritise therapeutically relevant genes. Accordingly, a wide range of network-based methods have been proposed to identify candidate therapeutic targets and reposition existing drugs, often leveraging metrics such as shortest path, nearest neighbour, and node centrality. For instance, centrality metrics have been used to highlight key hubs in disease-specific protein–protein interaction (PPI) networks [9, 10], while network proximity analyses have demonstrated strong predictive power in drug repurposing and target prioritisation tasks [7, 8, 11, 12].

Despite promising advances, most existing methods remain drug-centric, inherently limited by the scope of known compound-target interactions. Moreover, many frameworks operate on a single data modality, overlooking the available wealth of multi-omic and functional genomic data. Here, we introduce a multimodal machine learning framework that integrates single-cell RNA sequencing, bulk-omics, genome-wide CRISPR knockout data, and PPI networks to systematically profile and prioritise potential therapeutic targets in a disease-specific manner. Starting from publicly available datasets, we define disease-associated gene signatures, map them onto the human interactome, and extract network-informed descriptors for each gene. These features are then used to train off-the-shelf machine learning classifiers to distinguish known drug targets from non-targets, enabling systematic prioritisation of context-specific therapeutic candidates.

We applied this framework to clear cell renal cell carcinoma (ccRCC), the predominant subtype of kidney cancer, which accounts for approximately 80% of cases in a cancer that ranks sixth in prevalence in the UK [13]. It arises from proximal tubular epithelial cells (PTECs), with inactivation of the *VHL* tumour suppressor gene on chromosome 3p25 representing a hallmark initiating event [14–16]. In healthy PTECs, *VHL* encodes a component of an E3 ubiquitin ligase complex that targets hypoxia-inducible factors (HIF-1α and HIF-2α) for degradation. Loss of *VHL* leads to HIF accumulation under normoxic conditions, driving metabolic reprogramming, angiogenesis, and immune infiltration—features that collectively underpin the histopathological and molecular phenotype of ccRCC. Although tyrosine kinase inhibitors targeting the VEGF pathway and immune checkpoint inhibitors offer clinical benefit, a substantial proportion of patients exhibit intrinsic or acquired resistance, underscoring the need for alternative therapeutic strategies [17]. Notably, few existing RCC treatments act directly on tumour-intrinsic pathways, with the exception of HIF pathway inhibitors such as belzutifan, suggesting additional cell-intrinsic vulnerabilities remain unexploited. Using our framework, we identified and experimentally validated five candidates—ENO2, LRRK2, SCARB1, HMOX1, and TGM2—whose inhibition significantly impaired tumour cell viability, highlighting their potential as therapeutic targets.

## Results

### Identification of major cell populations in ccRCC by single-cell RNA sequencing

To characterise the cellular landscape of ccRCC at high resolution, we analysed single-cell RNA sequencing (scRNA-seq) data from 12 patients in the Li et al. cohort [18]. In that study, multiple anatomical regions were sampled from each donor for transcriptomic profiling, including the tumour core, tumour-normal interface, adjacent normal kidney, peripheral blood, perinephric fat, normal adrenal gland, adrenal metastasis, and tumour thrombus, where available. In the original study, transcriptomic data from all 12 donors were included in the downstream analyses. In contrast, our analysis focused exclusively on the 10 donors with histopathologically confirmed ccRCC, excluding the two non-RCC samples prior to re-clustering. Following quality control, we retained the transcriptomes of 250,331 cells, which were clustered to resolve 16 distinct broad cell types based on canonical marker gene expression and subsequently assessed for their distribution across tissue sites (**Fig. 1A-D**). Tumour cells were annotated based on CA9 over-expression, a marker absent in healthy renal tissue but strongly induced in ccRCC as a result of HIF-1α accumulation [19].

**Figure 1:**
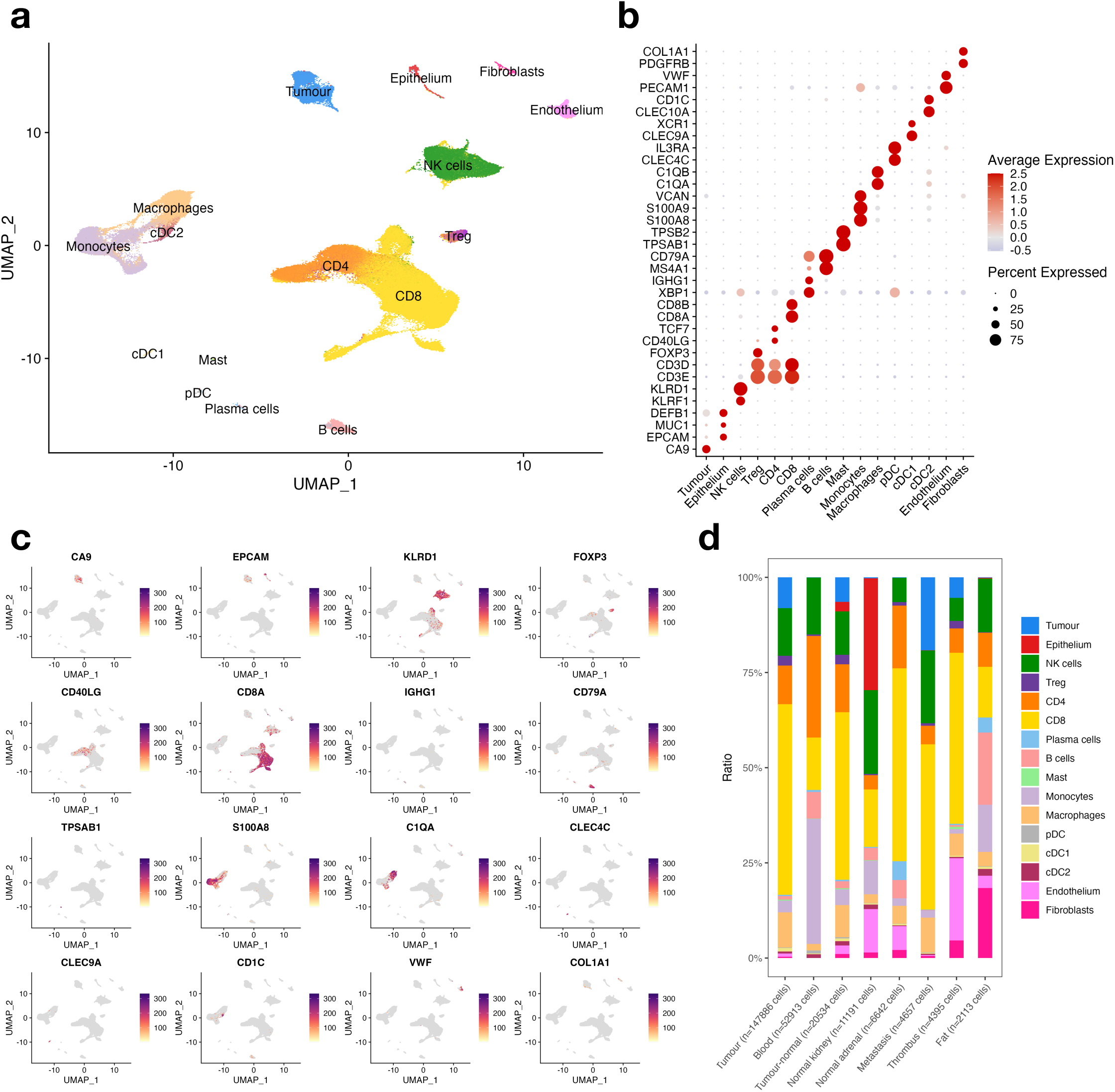
single-cell transcriptomic profiling of ccRCC reveals 16 broad cell compartments. **a** Uniform manifold approximation and projection (UMAP) visualisation of 250,331 cells obtained from 10 histopathologically confirmed ccRCC donors in the Li et al. cohort. Cells are coloured by the 16 broad cell compartments identified through unsupervised clustering and canonical marker gene expression. Multiple anatomical sites were sampled from each donor, including the tumour core, tumour-normal interface, adjacent normal kidney, peripheral blood, perinephric fat, normal adrenal gland, adrenal metastasis, and tumour thrombus, where available. **b** Expression of canonical cell type marker genes used to annotate the 16 cell compartments. **c** UMAP visualisation from (**a**), overlaid with the expression of representative marker genes used to define major cellular compartments. **d** Bar plot showing the distribution of the 250,331 profiled cells across eight anatomical regions sampled for sequencing.

### Transcription factor activity-based sub-clustering of epithelial and tumour cells

To achieve a more refined characterisation of the identified broad cell types, we performed sub-clustering analyses within each major population. A key limitation of current scRNA-seq technologies is the high gene drop-out rate, where transcripts from lowly expressed genes are often missed, leading to zero or near-zero read counts. We hypothesised that clustering cells based on inferred transcription factor (TF) activity, rather than directly using UMI data, could help mitigate this issue and provide a more informative broad cell type sub-clustering output. First, TFs are major determinants of a cell’s phenotype, and so their activity level can provide significant insights into the cell type of a particular cluster. Second, tools such as SCENIC [20] work as biological dimensionality reduction methods by aggregating the expression levels of a TF and its predicted targets into a single score. Consequently, regulon (defined as a TF and all its predicted targets) activity inference and its subsequent use in scRNA-seq clustering enables the identification of cell clusters based on groups of genes with functional relationships. Therefore, even if some genes in a regulon are not captured during sequencing due to drop-out, the presence of other genes in the same regulon allows for more accurate cell identity assignment.

Such TF-based sub-clustering of epithelial cells revealed 10 cell groups (**Fig. 2A-D**). These included two proximal tubule (PT) populations, consistent with the predicted HNF4A [21], HNF4G [22], and MAF [23] TF activities (**Fig. 2B**). Cluster 1 was distinguished by the highest expression of markers characterising all three PT segments [24] and is hypothesised to represent the cell of origin of ccRCC, while cluster 2 exhibited heightened expression of metallothionein family genes (**Fig. 2C**). Meanwhile, cluster 3 represents the loop of Henley (LoH), based on elevated levels of genes such as CLDN3, CLDN16 [25], and SLC12A1 [26]. Clusters 4 & 5 expressed markers of the distal convoluted tubule (DCT), examples including WNK1, SLC12A3, and TFAP2A [27] regulon activity (**Fig. 2B-C**), with cluster 5 also over-expressing immediate-early genes. Furthermore, two clusters were identified as collecting duct (CD) cells, with one corresponding to principal cells (AQP2, GATA2 & 3 [28]) and another to intercalated cells (ATP6V0D2, FOXI1 [29]). Cluster 8 was annotated as a podocyte population given the elevated PODXL gene expression (**Fig. 2C**) and up-regulated WT1 regulon activity [30] (**Fig. 2B**). The pelvic urothelial cluster was annotated based on AQP3 and S100P expression, consistent with the increased activity of TP63 TF [31], among others. Lastly, a cluster exhibiting elevated levels of genes involved in oxidative phosphorylation was observed, agreeing with the upregulated ATF5 [32] regulon activity.

**Figure 2:**
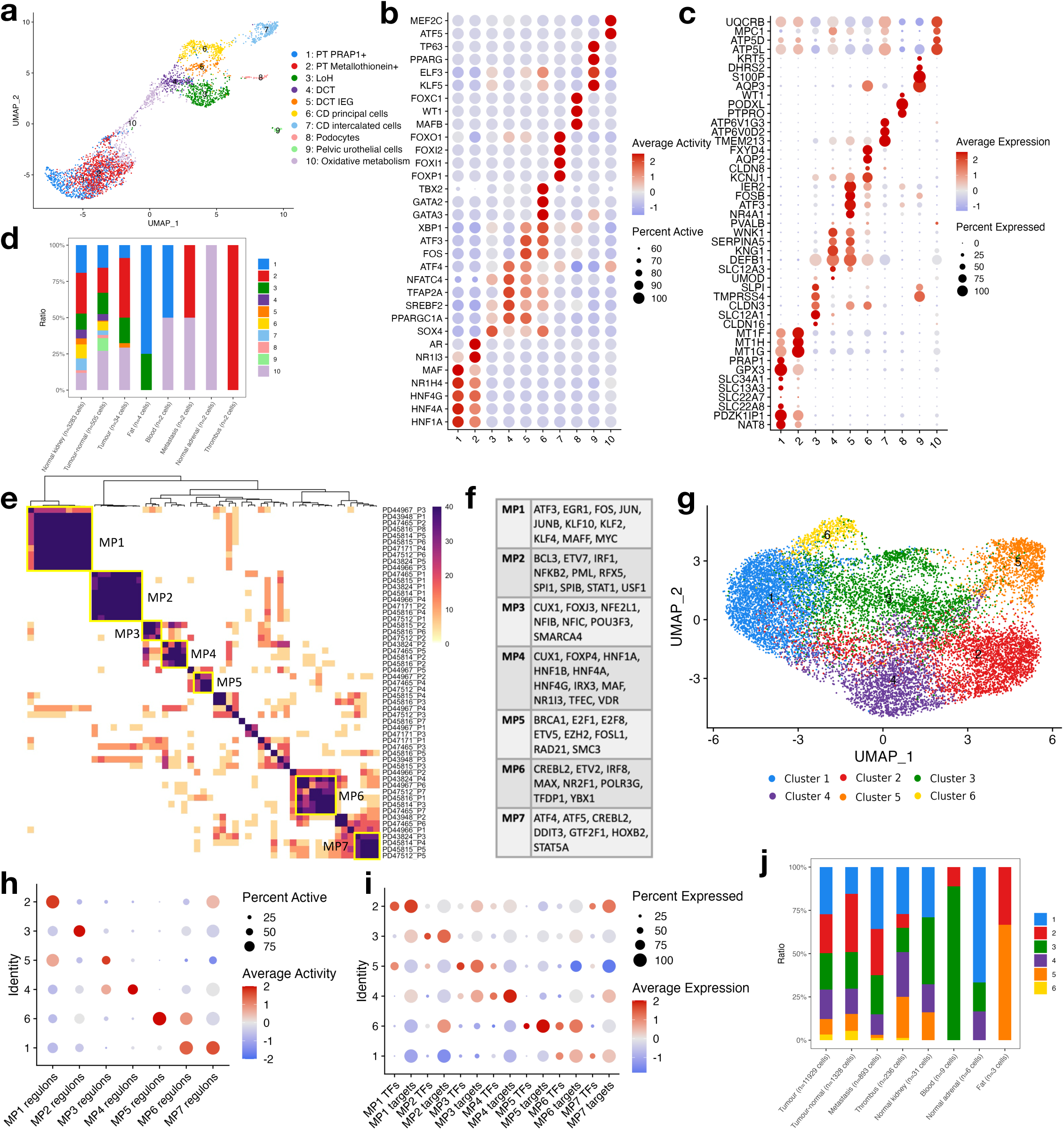
Transcription factor activity-based sub-clustering refines epithelial and tumour cell populations. **a** UMAP visualisation of epithelial cells following sub-clustering based on inferred transcription factor (TF) activities. Per-cell TF activity scores were computed using the SCENIC pipeline, and the resulting regulon activity matrix was used for clustering. Cluster identities were subsequently mapped back onto the scRNA-seq data to assess marker gene expression and validate annotations. **b** Inferred regulon (TF activity) scores across epithelial sub-clusters. **c** Expression of canonical epithelial marker genes across the identified sub-clusters. **d** Distribution of epithelial sub-clusters across anatomical regions sampled for sequencing. **e** Heatmap of 55 tumour transcriptional programs identified using non-negative matrix factorisation (NMF). Rows and columns represent TF activity programs across individual patients, with colouring intensity reflecting similarity between programs. Yellow squares indicate seven transcriptional meta-programs (MPs) shared across multiple patients. **f** Defining TFs for each of the seven identified tumour meta-programs. **g** UMAP representation of tumour cells following TF activity-based sub-clustering. **h** Average meta-program regulon activity across the identified tumour sub-clusters. **i** Average expression of representative TFs and their target genes defining each meta-program. **j** Distribution of tumour sub-clusters across anatomical regions sampled for sequencing.

In contrast, tumour cell analysis presents a greater challenge due to the uncertainty surrounding the number and characteristics of clusters that one should expect to see. To address this, we applied nonnegative matrix factorisation (NMF) to the TF activity profiles of tumour cells from for each patient individually [18, 33]. This approach facilitated the identification of transcriptional programs that are shared by multiple patients, thus representing transcriptional ‘meta-programs’ (MPs). Overall, the outlined analysis of tumour cells yielded a total of 55 transcriptional programs and 7 transcriptional meta-programs (**Fig. 2E-F**), which were then assessed for Reactome [34] pathway enrichment (**Supplementary Fig. 1**). MP1 showed enrichment in mRNA translation pathways, while MP2 was distinguished by a significant presence of immune interaction pathways. In contrast, MP3 was linked to a single pathway, namely FOXO-mediated transcription. MP4 displayed enrichment in pathways typically found in tubular kidney cells, including SLC-mediated transmembrane transport and lipid metabolism. MP5 encompassed cell division pathways, while MP6 was associated with mRNA translation, cell division, as well as several immune pathways. Finally, MP7 was characterised by pathways that are typically upregulated during starvation. Once meta-programs were established, tumour cells were clustered. As expected, different clusters associated with different meta-programs, with a given cluster expressing at most 2 MPs (**Fig. 2G-J**).

The outlined TF-based sub-clustering was subsequently applied to the remaining broad cell types (**Supplementary Fig. 2, Supplementary Tables 1 & 2**). Overall, the approach proved to be advantageous, enhancing cluster resolution by partially overcoming the limitations imposed by gene drop-out in scRNA-seq data and providing additional insights in the form of inferred TF activities. For instance, within the CD4+ T cell compartment, it enabled clear distinction of Th1, Tfh, and Th17 helper cell subsets—populations that were not resolved in the original study due to the near-complete dropout of both key transcription factors and conventional surface markers typically used to define these subsets.

### Network analysis for characterisation of candidate disease-associated genes

To gain a comprehensive understanding of protein-coding gene functions in relation to ccRCC, we compiled 34 candidate disease-associated gene (CDAG) signatures as well as a list of ccRCC drug target genes (DTGs; **Supplementary Table 3**), outlined in **Table 1**. The CDAG sets include scRNA-seq gene expression and TF activity differences between tumour cells and epithelial cluster 1 (PT PRAP1+ cells), scRNA-seq broad cell type marker genes, differentially expressed genes, proteins and phosphoproteins obtained from bulk ccRCC omics data analyses of tumour versus healthy tissues, most commonly lost or gained genes identified in ccRCC whole-genome sequencing data, and ccRCC-related signatures from DisGeNET [35], the Human Protein Atlas [36], and DepMap [37] databases. Meanwhile, the DTG set comprised 202 genes targeted by FDA-approved ccRCC treatments (25 genes) as well as genes targeted by drugs in phase I-IV ccRCC clinical trials (177 genes). Recognising that genes within biological systems reveal their functions when viewed in relation to one another, we mapped the DTG and CDAG signatures onto the human protein-protein interaction network (PPI; STRING [38] database, confidence threshold ≥400; 19,303 proteins) and interrogated their average centralities (**Fig. 3A**). This analysis revealed that DTGs exhibit the highest average degree, eigenvector, and PageRank centralities, underscoring their roles as hub nodes. Next, we characterized the DTG set in relation to the compiled 35 signatures by comparing the average shortest path distances within the PPI network (**Fig. 3B, Supplementary Fig. 3**). Specifically, we calculated the average shortest paths from each gene in the DTG set to every gene in a given signature and compared them to the average shortest paths from each DTG to all other nodes in the network not part of that signature. Overall, DTGs were found to be closer to the curated gene sets than to the rest of the gene population, with the exception of downregulated bulk RNA-seq transcripts, genes with copy number gains, and those classified under the “Others” category.

**Figure 3:**
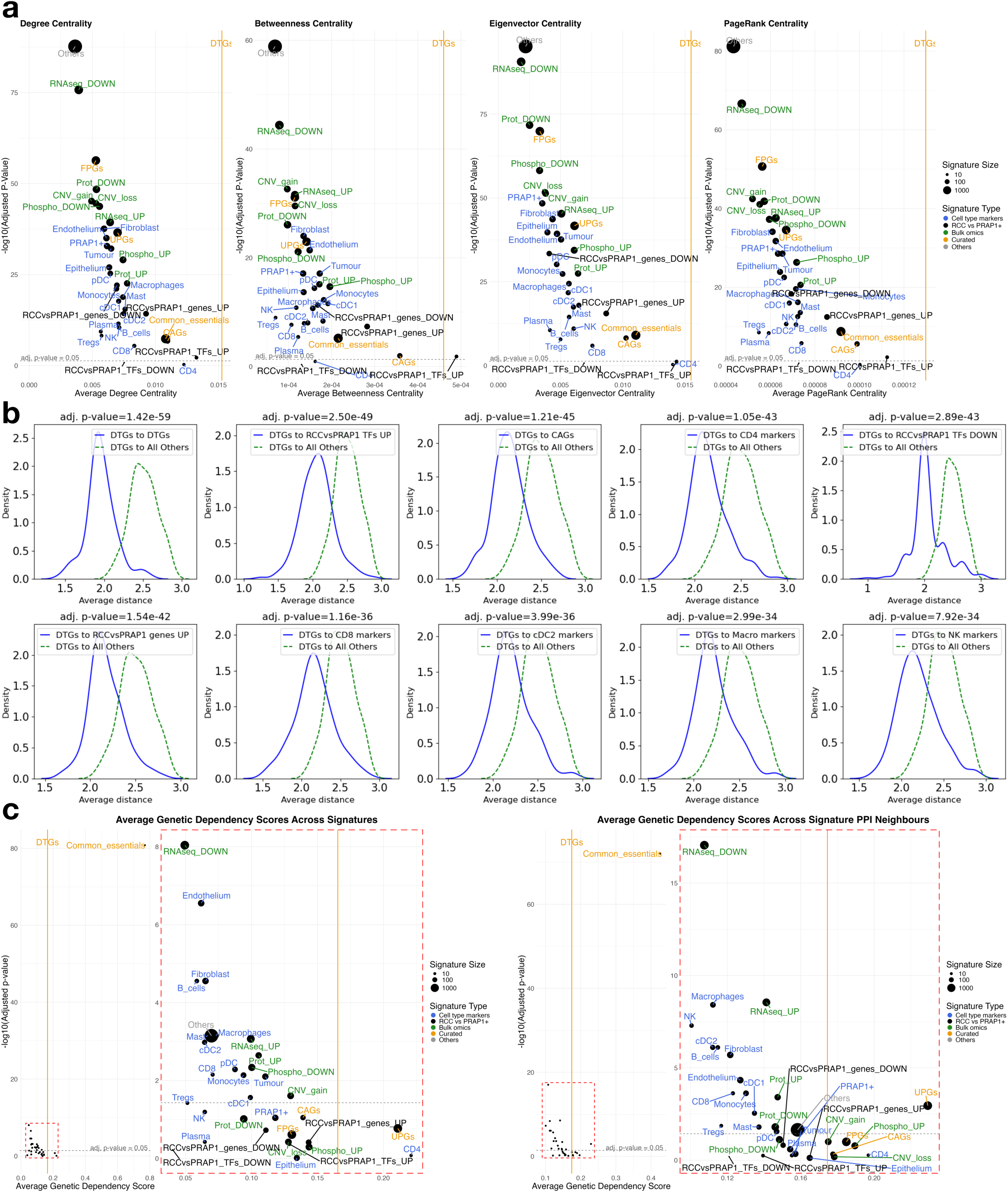
Drug target gene and candidate disease associated gene signature characteristics. **a** Relationship between average network centrality and statistical significance of centrality differences relative to DTGs. The *x*-axis shows the average centrality score (degree, betweenness, eigenvector, or PageRank) of each signature within the human protein–protein interaction (PPI) network. The *y*-axis represents the statistical significance of the difference in centrality between each CDAG set and the DTG set, expressed as –log₁₀ of the BH-adjusted *p*-value (Wilcoxon rank-sum test). **b** Comparison of average shortest path distance distributions within the PPI network, measuring the proximity of DTGs to a given signature (blue) versus to all remaining network nodes (green). Top 10 most significantly different distributions are shown. Statistical significance was assessed using BH-adjusted Wilcoxon rank-sum tests; adjusted *p*-values are indicated. **c** Average genetic dependency of CDAG signatures relative to DTGs, based on CRISPR knockout screens across 16 ccRCC cell lines (DepMap data). The left panel shows results for genes within each signature; the right plot shows results for the immediate PPI neighbours of each signature. The *x*-axis indicates the average dependency scores, and the *y*-axis corresponds to –log₁₀ of the BH-adjusted *p*-value for dependency differences (Wilcoxon rank-sum test).

**Table 1:**
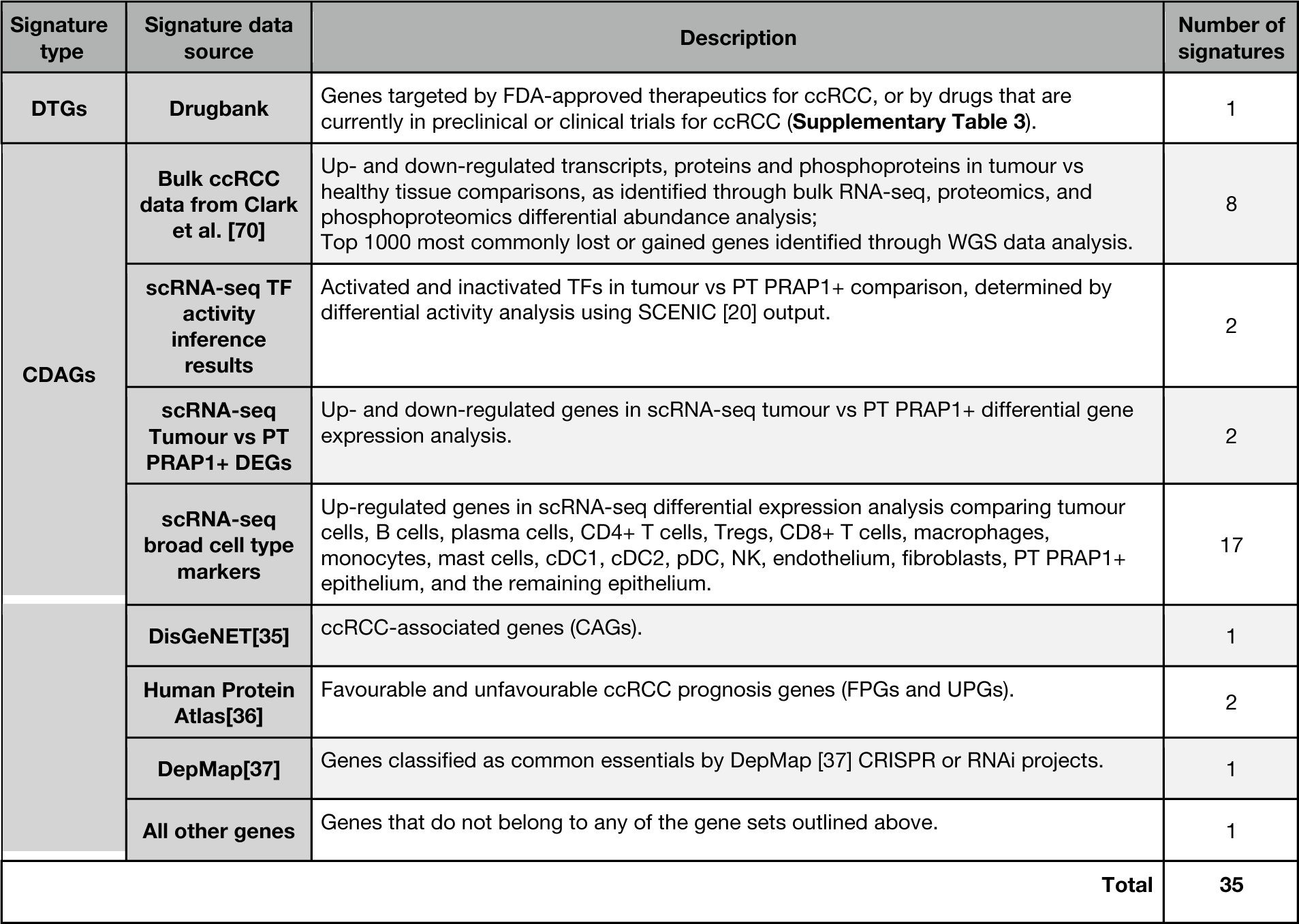
Drug target and candidate disease-associated gene signature descriptions.

We also investigated gene essentiality across 16 ccRCC cell lines using systematic CRISPR knockout data from the DepMap [37] database, focusing on 17,880 proteins shared between the STRING PPI network and DepMap data. Dependency scores ranging near 1 signify gene’s criticality for cell survival—its knockout leading to marked growth inhibition or cell death. Conversely, scores near 0 imply negligible impact on cell growth upon gene knockout. Notably, dependency data was not available for three DTGs (CSF2RA, IL3RA, and XRCC7). A comparative analysis of the average gene dependency scores between the remaining 199 DTGs and CDAGs revealed that only one gene set, the common essential genes, exhibited significantly higher dependency scores. (**Fig. 3C**). This observation is expected given that common essential genes are defined as the most consistently depleted across multiple cell lines. The lack of significantly higher average dependency in any other gene set indicates that DTGs tend to cause more pronounced growth arrest in ccRCC cell lines. However, it is important to note that DTGs are not universally essential for ccRCC lines, as genes are considered essential only if their dependency score exceeds 0.5, whereas the average for DTGs is ∼0.17. This observation aligns with the fact that most ccRCC therapeutics directly affect the tumour microenvironment rather than the tumour cells themselves, as seen with VEGF pathway inhibitors and immunotherapies. A similar trend was observed when comparing the average dependency scores of each gene set’s PPI neighbours, with the common essential gene neighbourhood again showing the highest average dependency.

Overall, the outlined results confirm that the curated signatures were not merely random selections of genes but are functionally relevant to the disease. Namely, the relatively high DTG network centralities indicate their involvement in multiple pathways, which, when disrupted, lead to disease regression. This finding is further supported by the comparatively high genetic dependency scores associated with DTGs. Finally, the close proximity among DTGs suggests that they form a network module, while their proximity to CDAGs indicates that CDAGs are involved in pathways essential for cancer maintenance and development.

### Machine learning model construction and performance

Our findings align with the work of Isik et al. [11], who demonstrated that the proximity between drug targets and genes deregulated by the same drug yields the highest target prediction accuracy. Building on this insight, we extended and generalised the approach by leveraging 35 disease signatures to compute 139 features for each of the 17,880 genes, capturing both their genetic dependency profiles and their positioning within the PPI network (**Supplementary Table 4**). Therefore, instead of predicting targets of a particular compound based on its perturbation profile, such gene-level characterisation allows us to predict the so-called targets of the disease given multiple disease signatures. These 139 features were subdivided into six categories (**Fig. 4A**): genetic dependency scores across 16 ccRCC cell lines (16 features); average genetic dependency scores of a gene’s PPI neighbours across the same 16 ccRCC cell lines (16 features); PPI network centralities (4 features); proportions of a gene’s PPI neighbours that belong to CDAG or DTG signatures (35 features); average shortest path length from a given gene to all genes in a particular CDAG or DTG signature (35 features); and finally, minimum shortest path distance from a given gene to a member of a particular CDAG or DTG signature (33 features; minimum distances to DTGs and genes belonging to the ‘Others’ set were excluded to avoid label leakage and prevent biasing predictions towards already known gene-disease associations).

**Figure 4:**
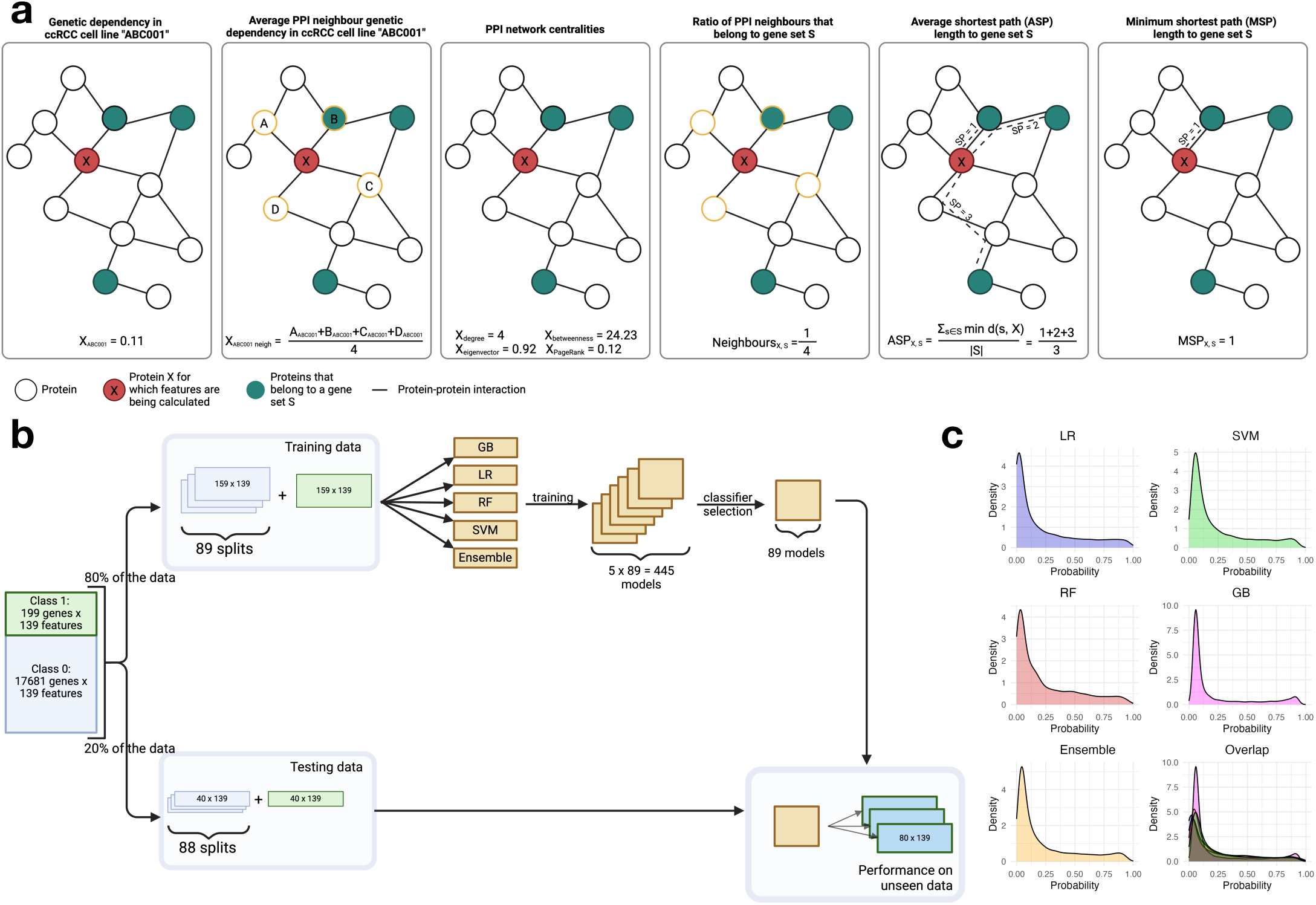
Machine learning for the identification of novel therapeutic targets for ccRCC. **a** 6 types of features used for gene characterisation. Genetic dependency data across 16 ccRCC cell lines, obtained from the DepMap database, provided 16 features representing the gene’s dependency score in each cell line, along with another 16 features capturing the average dependency scores of the gene’s PPI neighbours in those same cell lines. Network-based features include degree, betweenness, eigenvector, and PageRank centralities, yielding 4 features per gene. Additionally, the 35 curated disease signatures were used to compute: (i) the proportion of a gene’s PPI neighbours belonging to each gene set (35 features), (ii) the average shortest path length to each gene set (35 features), and (iii) the minimum shortest path length to each gene set (33 features). **b** Machine learning training and testing workflow. Protein-coding genes were split into two classes: Class 1, comprising ccRCC drug target genes, and Class 0, encompassing all other genes. Subsequently, an 80/20 split partitioned these classes for model training and testing purposes. To ensure balanced representation during training and testing, Class 0 genes were randomly sampled exactly once to match the count of Class 1 genes. Each resulting balanced dataset was then used to train four machine learning classifiers and their ensemble to distinguish the two classes of genes. Classifier performance was assessed via five-fold cross-validation, and the overall best-performing method was selected for evaluation on the held-out test set and for final predictions across the full dataset. **c** Class 1 probability density distributions across the applied ML classifiers.

Building on the work by Tsagkogeorga et al. [39], we next designated genes encoding DTGs as the positive class (Class 1) and all other genes as the negative class (Class 0), with the aim of developing a machine learning predictor capable of distinguishing these two classes of genes using the 139 features. We partitioned these classes in an 80/20 split for model training and testing (**Fig. 4B**). To ensure balanced datasets, negative class genes in both training and testing sets were randomly sampled exactly once to match the number of Class 1 genes. Subsequently, four machine learning classifiers (Logistic Regression (LR), Support Vector Machine (SVM), Random Forest (RF), and Gradient Boosting (GB)) and their ensemble were trained using the 89 balanced training datasets. Evaluation metrics, derived from 5-fold cross-validation (CV), are presented in **Table 2**. The ensemble classifier outperformed the remaining ML methods in all metrics except precision.

**Table 2:** Performance metrics across 5 ML classifiers obtained using 5-fold CV on the training data.

|  | Ensemble | GB | LR | RF | SVM |
| --- | --- | --- | --- | --- | --- |
| Accuracy | 0.8475 ± 0.0298 | 0.8461 ± 0.034 | 0.8313 ± 0.0352 | 0.8409 ± 0.0363 | 0.8393 ± 0.0326 |
| Precision | 0.8361 ± 0.0361 | 0.837 ± 0.0427 | 0.8189 ± 0.0428 | 0.8326 ± 0.0426 | 0.8257 ± 0.0406 |
| Recall | 0.8688 ± 0.0341 | 0.8647 ± 0.0408 | 0.8558 ± 0.0342 | 0.8581 ± 0.0466 | 0.865 ± 0.0546 |
| F1 | 0.8504 ± 0.0283 | 0.8487 ± 0.0327 | 0.835 ± 0.033 | 0.8432 ± 0.0359 | 0.8429 ± 0.0332 |
| AUC | 0.9173 ± 0.025 | 0.9147 ± 0.0264 | 0.8993 ± 0.0312 | 0.9151 ± 0.0258 | 0.9068 ± 0.0287 |

We next examined feature importance estimates to gain insight into the factors driving model predictions (**Supplementary Fig. 4**). Among the top-ranked features were the average shortest path distances to immune cell marker signatures, as well as the neighbourhood ratio and average shortest path to the DTG set. Notably, no single feature dominated the importance scores, suggesting that predictive performance derives from the combined contribution of multiple features rather than being driven by any one in isolation. Regarding probability distributions across all 17,880 assessed genes as predicted by the five classifiers, a high degree of similarity was observed, evidenced by their substantial overlap (**Fig. 4C**). Most genes were assigned a probability near zero, aligning with the hypothesis that the majority of genes are unlikely to be viable drug targets. Finally, the strong agreement among the top 1% of predictions across classifiers further supports result reliability (**Supplementary Fig. 5**).

### New therapeutic target prediction

Based on the superior performance of the ensemble machine learning method, we selected it for the prediction of novel drug targets and their subsequent validation in ccRCC cell lines (**Supplementary Table 5**). The ensemble model exhibited the following performance metrics on the hold-out data (40 DTGs and 3,530 negative class genes): accuracy of 0.8569, precision of 0.8335, recall of 0.900, F1 score of 0.8644, and AUC of 0.9254. To estimate the false positive rate, average Class 1 probabilities were calculated for the hold-out data across the 89 ensemble models. The number of Class 0 genes that were incorrectly assigned a Class 1 probability above 0.5 was 660 out of 3,530, resulting in the false positive rate of 0.187.

To identify potential novel therapeutic targets, we utilised AstraZeneca’s 5 R’s R&D [40] framework (right target, right patient, right tissue, right safety, right commercial potential). For the ‘Right target’ and ‘Right tissue’ criteria, we focused on the top 10 novel predictions that directly impact tumour cells. This selection process considered genes markedly over-expressed in bulk proteomic data or, if such data were unavailable, transcriptomic data from tumour versus normal tissue comparisons (log2FC > 0.5, adj. p-value < 0.05). Candidate genes were further required to be overexpressed in tumour cells relative to all other broad cell types in scRNA-seq data (log2FC > 0.25, adj. p-value < 0.05), as well as relative to the PT PRAP1+ epithelial cluster (log2FC > 0.5, adj. p-value < 0.05). In addition, genes were filtered for high cell type specificity, defined by a scRNA-seq Tau score greater than 0.5 as reported by the Human Protein Atlas. The top 10 candidates that fit the criteria were: Lysyl oxidase (LOX), Enolase 2 (ENO2), Leucine rich repeat kinase 2 (LRRK2), Transglutaminase 2 (TGM2), Heme oxygenase 1 (HMOX1), Scavenger receptor class B member 1 (SCARB1), Caveolin-1 (CAV1), Transforming growth factor alpha (TGFA), c-Myc (MYC), and Intercellular Adhesion Molecule 1 (ICAM1).

For ‘Right safety,’ genes identified as common essentials by DepMap [37] CRISPR or RNAi projects, namely MYC, were excluded. The ‘Right patient’ criterion was met by focusing exclusively on ccRCC data, and the ‘Right commercial’ aspect was supported by the high prevalence of kidney cancer and the unmet clinical need in patients who experience primary progression despite current standard-of-care therapies. The remaining nine genes were further narrowed down by removing those without available small-molecule inhibitors, resulting in the exclusion of CAV1 and TGFA. Finally, we also excluded immune-related predictions—ICAM1—in order to permit validation of our hypotheses in a non-immune experimental model. Visualisation of the ML feature space revealed a highly cohesive clustering of known and newly identified targets (**Fig. 5A**). This, in turn, further demonstrates that the 139 features used to characterise each gene capture the biological information that determines whether that gene is a likely ccRCC therapeutic target.

**Figure 5:**
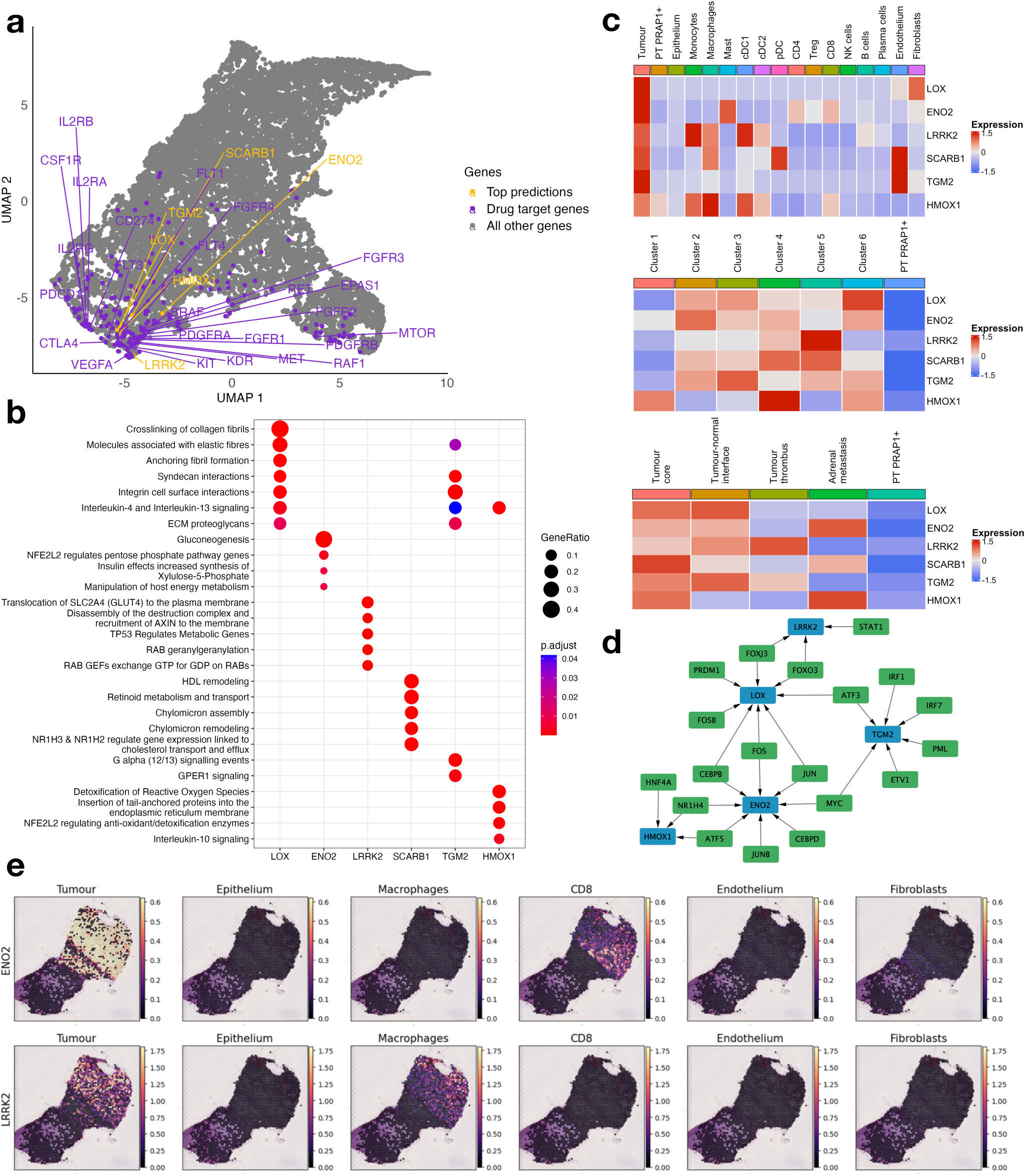
Characterisation of top tumour target predictions. **a** UMAP visualisation of the feature space used to train and evaluate the machine learning target prediction framework, comprising 17,880 genes and 139 features. Genes belonging to the drug target gene (DTG) set are highlighted in purple. Within this group, genes targeted by FDA-approved therapies for ccRCC are highlighted and labelled in purple, while those currently under investigation in clinical trials are highlighted in purple without labels. The refined set of top tumour target predictions is highlighted in yellow and labelled. **b** Reactome pathway enrichment analysis of the protein-protein interaction (PPI) neighbourhoods of the six top tumour target predictions. High-confidence interaction partners (confidence score ≥700) were retrieved from the STRING database to define each candidate’s PPI neighbourhood. Reactome pathway enrichment was performed on these neighbourhoods, and results were filtered to retain only pathways corresponding to terminal nodes (leaf pathways) in the Reactome hierarchy, ensuring specificity of functional annotations. **c** scRNA-seq expression profiles of the six top candidate genes. Top: expression levels across broad cell types. Middle: expression patterns across tumour and proximal tubule PRAP1⁺ (PT PRAP1⁺) cell clusters. Bottom: expression comparison between tumour and PT PRAP1⁺ cells, stratified by the anatomical region sampled for sequencing. **d** Inferred transcriptional regulators of the six predicted targets in tumour cells, identified using the SCENIC pipeline. **e** ENO2 and LRRK2 expression patterns across indicated cell types at the tumour-normal interface in spatial transcriptomics data from ccRCC patient PD47171.

To elucidate the functions of the remaining predicted targets, Reactome [34] pathway enrichment analysis was conducted for the high confidence PPI neighbourhoods of these genes (**Fig. 5B**; confidence score ≥700). The results revealed that LOX and TGM2 are involved in extracellular matrix organisation, ENO2 is associated with glucose metabolism, LRRK2 participates in multiple pathways including GLUT4 translocation and membrane trafficking, HMOX1 is involved in the inactivation of reactive oxygen species, and SCARB1 plays a role in lipid metabolism. Regarding target scRNA-seq expression profiles, all six genes demonstrated good tumour specificity, with the additional cell types expressing some of these genes being myeloid and endothelial cells (**Fig. 5C**). Next, SCENIC [20] output was analysed to identify target transcriptional regulators. SCARB1 had no predicted TFs regulating its expression, likely due to the stringent filtering criteria applied (see Materials & Methods section). The remaining five genes, along with their predicted TFs, formed a connected network, as depicted in **Figure 5D**. Finally, spatial transcriptomics data of the ccRCC tumour core and the tumour-normal interface were evaluated. The results corroborated scRNA-seq findings, confirming that the six target genes are specific to tumour rather than the adjacent normal tissues (**Fig. 5E; Supplementary Fig. 6**).

### Experimental validation of predicted targets

The six predicted targets were subsequently evaluated in three ccRCC cell lines using small-molecule inhibitors. Cytotoxicity assays revealed that inhibition of ENO2 (POMHEX[41], IC50=28.9nM) LRRK2 (LRRK2-IN-1 [42], IC50=13nM), and SCARB1 (BLT-1[43], IC50=50nM) produced the most pronounced reduction in cell viability, with the corresponding average inhibitor cytotoxicity IC50 values across the tested cell lines being 0.875, 22.9, and 55.7μM (**Fig. 6A, C; Supplementary Fig. 7**). This was further supported by proliferation assays (**Fig. 6B; Supplementary Fig. 8**), where treatment with 10μM of the respective inhibitors led to an average reduction in cell proliferation of approximately 45% for SCARB1 and LRRK2, and nearly complete inhibition for ENO2.

**Figure 6:**
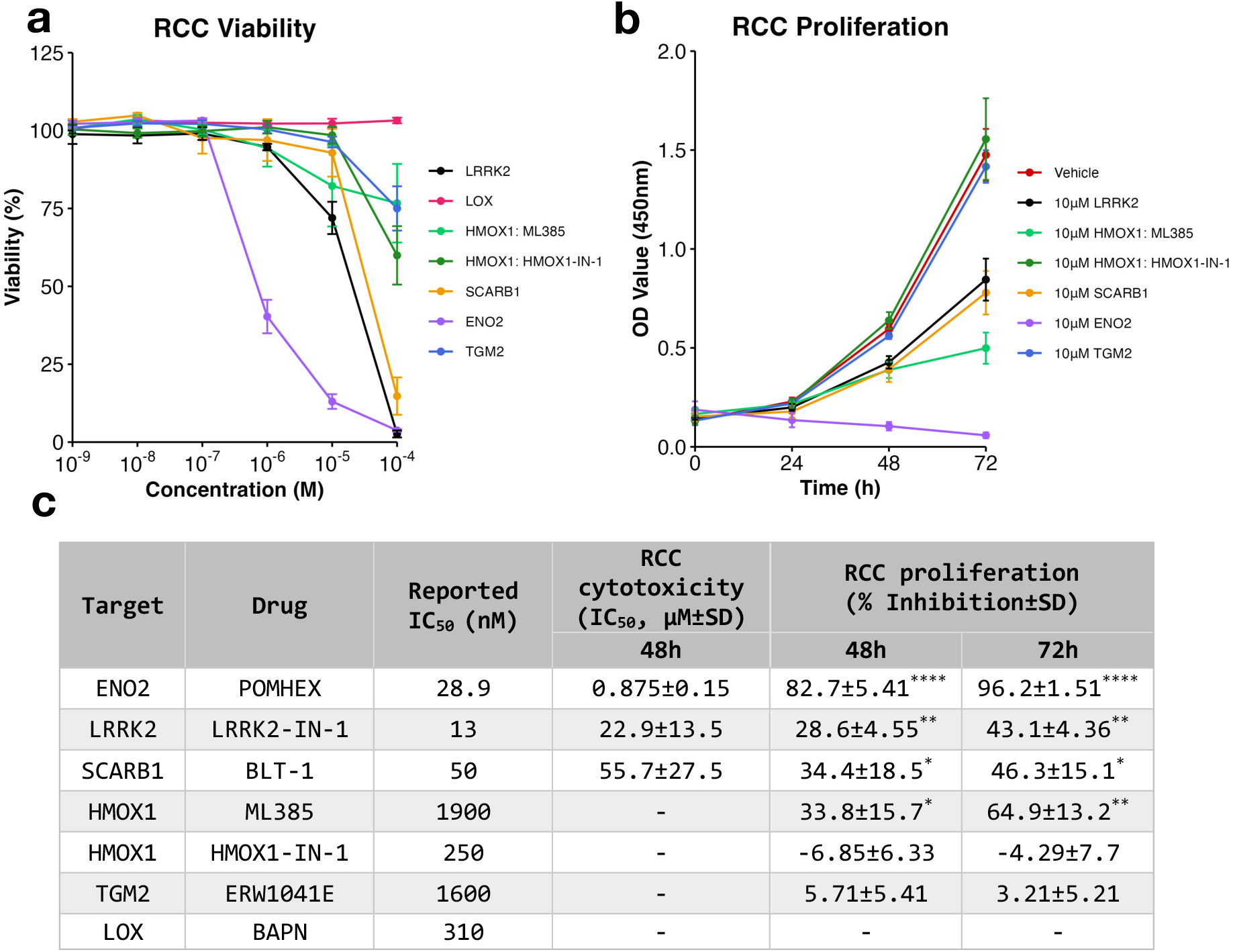
Experimental validation of top tumour target predictions. **a** Cytotoxicity assay results. Three ccRCC cell lines were treated in triplicate for 48 hours with either vehicle controls or inhibitors targeting LRRK2, LOX, HMOX1 (via its upstream regulator NFE2L2 using ML385 or directly using HMOX1-IN-1), SCARB1, ENO2, and TGM2, across six concentrations (1nM, 10nM, 100nM, 1µM, 10µM, 100µM). Cell viability was calculated relative to DMSO (10^-5^ or 10^-4^M; LRRK2, HMOX1, SCARB1, ENO2, TGM2) or ddH2O (10^-4^M; LOX) treated controls after background subtraction. For each cell line, technical replicates were averaged, and the resulting cell line means were used to calculate the overall mean±SEM across the three cell lines. **b** Proliferation inhibition assay results. Three ccRCC cell lines were treated in triplicate for 72 hours with either vehicle controls or 10µM of inhibitors targeting the genes of interest. Background-corrected absorbance values are reported as mean±SEM across three cell line averages. **c** Summary of drug response in RCC. Data are presented as mean±SD across three cell line averages. Asterisks indicate significance compared to vehicle control (Kruskal Wallis, Dunn’s multiple comparison; *P*<0.05*, P<0.01**, ****P<0.0001). Dashes indicate conditions that were not tested or for which values could not be determined.

For HMOX1, two drugs were tested: one directly targeting the gene (HMOX1-IN-1 [44], IC50=0.25μM), and one inhibiting the upstream regulator NFE2L2 [45] (ML385 [46], IC50=1.9μM). Along with TGM2 suppression (ERW1041E [47], IC50=1.6μM), inhibitor effects were cell line-specific: 786-O cells were most sensitive to ML385 and ERW1041E, whereas A498 cells showed the greatest sensitivity to HMOX1-IN-1. The comparatively weaker response to TGM2 and HMOX1 inhibition, relative to ENO2 or LRRK2, is likely attributable to the higher IC50 values of the respective inhibitors. Higher compound concentrations were not considered due to DMSO-associated cytotoxicity above 10^-4^M (**Supplementary Fig. 9**). In contrast, targeting LOX (β-Aminopropionitrile [48], IC50=0.31μM) did not affect RCC cell viability.

## Discussion

In this study, we have presented a multimodal framework that integrates disease-associated gene signatures, genome-wide CRISPR knockout data, and PPI network features to systematically prioritise therapeutic targets. Applied to clear cell renal cell carcinoma, this approach identified five candidate genes, among which ENO2 and LRRK2 demonstrated the most potent anti-ccRCC effects, highlighting their potential as targets for therapeutic intervention.

The key strength of the framework lies in its capacity to integrate data from multiple modalities, providing a flexible and scalable approach that can be readily extended to other diseases. By mapping disease signatures onto the human interactome and computing both average and minimum shortest path distances to each gene, we quantify how readily a particular node can communicate with the signatures. These topological features are further complemented by local PPI neighbourhood information and CRISPR knockout data, generating a feature space that characterises each gene with respect to the disease of interest. Crucially, this approach does not rely on known functional associations such as pathway data, thereby mitigating biases toward well-characterised biological processes. These features are then fed into a machine learning pipeline that addresses class imbalance and uses standard classifiers to discriminate known drug targets (Class 1 genes) from the broader gene pool. Beyond target prioritisation, the flexibility of the framework also allows it to be adapted to other biological questions—such as gene function inference—provided an appropriate positive class can be defined. As such, this method represents a scalable and generalisable strategy for extracting actionable insights from heterogeneous biological data.

Another key advantage of the approach is its ability to anchor predictions in empirical evidence. By training on known drug targets, the model learns not only from topological context but also from functional perturbation outcomes, providing a more biologically informed alternative to unsupervised approaches that rank genes solely based on individual topological features such as centrality or proximity [7, 12]. For example, while DTGs exhibit higher centrality and closer proximity to disease signatures, feature importance results revealed that no single feature was dominating. Instead, predictive performance arose from the combined contribution of multiple features, with shortest path-based metrics— particularly average distance to disease signatures—emerging as the most informative.

While the framework demonstrates strong predictive performance, several limitations should be acknowledged. First, although it effectively prioritises candidate targets, it does not provide mechanistic insights into how or why targeting a particular gene may confer therapeutic benefit. As such, downstream experimental validation remains essential to establish causal relevance. Second, the approach relies on protein–protein interaction networks, which remain incomplete and subject to biases arising from uneven research attention across genes. Third, the classification of genes into positive (known targets) and negative (non-targets) classes is inherently imperfect. In the context of ccRCC, many of the curated drugs remain under clinical investigation, and some genes labelled as non-targets may, in fact, represent emerging or as-yet-unvalidated therapeutic candidates. Nonetheless, the model’s high accuracy (0.8569) and AUC (0.9254) on unseen data support its practical utility for target prioritisation, even in the presence of data incompleteness and label uncertainty.

Given the limited availability of drugs that directly target ccRCC tumour cells rather than the tumour microenvironment, we focused on top-ranked candidates with tumour cell specificity. Functional validation across three ccRCC cell lines demonstrated that inhibition of ENO2 (POMHEX, IC50=28.9nM), LRRK2 (LRRK2-IN-1, IC50=13nM) and SCARB1 (BLT-1, IC50=50nM) produced the most potent and consistent effects. These results align with prior studies reporting that knockdown of either ENO2 [49] or SCARB1 [50] significantly suppresses ccRCC growth both in vitro and in vivo. Mechanistically, ENO2 knockdown induces ferroptosis - a nonapoptotic, iron- and lipid-dependent form of cell death [51]. Ferroptosis represents a targetable vulnerability in ccRCC, with prior studies demonstrating that HIF-2α is the central driver of this vulnerability through selective enrichment of polyunsaturated lipids, the rate-limiting substrates for lipid peroxidation [52, 53]. Similarly, SCARB1 is also heavily involved in lipid metabolism, with ccRCC relying on SCARB1 for high-density lipoprotein (HDL) import. Genome-wide association studies (GWAS) have identified a single nucleotide polymorphism at the SCARB1 locus associated with increased ccRCC risk, with Mendelian randomisation analyses of GWAS data further supporting a causal relationship between genetic alleles linked to elevated circulating HDL levels and ccRCC occurrence [54, 55].

For LRRK2, while its role has primarily been studied in the context of neurodegeneration— with its inhibitor BIIB122 currently undergoing phase III clinical trials for Parkinson’s disease—emerging evidence supports its relevance in ccRCC. Yang et al. [56] identified LRRK2 as a putative prognostic biomarker in ccRCC, reporting that aberrant LRRK2 expression is associated with altered DNA methylation patterns and that LRRK2 knockdown significantly reduces A498 and 786-O cell line proliferation. In parallel, GWAS results have linked LRRK2 to elevated blood cholesterol levels [57, 58]. Based on these findings, and prior to the publication of Hong et al. [59], we selected LRRK2 for further investigation, as its role in ccRCC had, to our knowledge, been explored only once previously - in the study by Yang et al. [56]. Hong et al. [59] independently demonstrated that LRRK2 knockdown in vivo significantly slowed ccRCC tumour progression and that LRRK2 promotes resistance to tyrosine kinase inhibitors and immunotherapy by stabilising a lipid metabolism gene LPCAT1. We note that this study was published after our model predictions had been finalised, highlighting that our machine learning framework can uncover clinically meaningful therapeutic targets independently of prior experimental reports.

In contrast, responses to HMOX1 and TGM2 inhibition were more variable and appeared to be cell line-dependent, although the comparatively lower potency of their respective inhibitors limits direct comparison with the other compounds. Previous studies have shown that HGF-MET signalling induces the Ras-Raf-ERK pathway, which, in turn, activates HMOX1 to counteract oxidative stress. Consistently, ccRCC cells treated with HGF show increased NFE2L2 nuclear translocation compared with vehicle-treated controls [60]. Furthermore, inhibition of HMOX1, either alone or in combination with MET, has been reported to reduce ccRCC tumour volume, decrease tumour vessel density, and increase oxidative stress in vivo [61]. Finally, HMOX1 downregulation is also known to cause ccRCC mitotic delay at the G2/M phase [62]. Our findings support these observations, with NFE2L2 inhibition resulting in an average proliferation reduction of 65% across the three cell lines. Meanwhile, TGM2 blockade has been shown to induce apoptosis in ccRCC mouse xenograft models through p53 stabilisation [63–65]. The resulting elevated p53 levels were also found to reduce angiogenesis by decreasing HIF-1α activity through the promotion of the p300–p53 interaction and the concurrent reduction in the p300–HIF-1α interaction [66].

In summary, this study presents a novel computational framework for systematic therapeutic target discovery, integrating disease signatures, functional genomics, and PPI network analysis within a machine learning pipeline. The identification of multiple ccRCC-relevant candidates, including ENO2 and LRRK2, highlights the effectiveness of the approach in uncovering functionally meaningful targets. The framework’s flexibility and scalability allow it to be applied to other diseases, providing a structured approach to target prioritisation in diverse biological contexts.

## Acknowledgements

The authors thank Dr. Ruoyan Li for assistance with data analysis during the early stages of this work.

## Contributions

G.B. conducted data analysis. I.T. carried out experimental validation. J.J. advised on the translational potential of candidate targets from a clinical perspective. N.H. and K.S.P. provided overall guidance.

## Materials & Methods

### scRNA-seq data acquisition and pre-processing

The scRNA-seq dataset was downloaded from Mendeley Data (https://data.mendeley.com/datasets/g67bkbnhhg/1). The data was subset to exclude two patients (PD44714 and PD47172) who had malignancies other than ccRCC.

### Broad cell type identification

The filtered Seurat [67] object was split according to the patient of origin using the *SplitObject* function, followed by normalisation of UMI counts (*NormalizeData*) and identification of 2000 most variable features (*FindVariableFeatures*) for each patient independently. Next, features that exhibited repeated variation across patients were determined using the *SelectIntegrationFeatures* function, and principal component analysis (PCA) was performed separately for each patient using these features. Integration anchors were subsequently calculated using the *FindIntegrationAnchors* function, setting the reduction parameter to ‘rpca’, followed by dataset integration (*IntegrateData*). Finally, the standard workflow involving data scaling (*ScaleData*), PCA calculation (*RunPCA*), UMAP drawing (*RunUMAP*), nearest-neighbour graph construction (*FindNeighbors*), and cluster determination (*FindClusters*) was carried out. Differentially expressed genes among the broad cell types were determined using the *FindAllMarkers* function; DEGs with log2FC > 1 and adjusted p-value < 0.05 were used to define the broad cell type marker gene lists.

### TF activity inference

The level of TF activity was estimated using pySCENIC [20] (v.0.9.15; Python implementation of SCENIC). Due to the exceptionally large number of identified cells, it was not computationally feasible to perform single-cell regulatory network calculation for the entire dataset in a single run. To circumvent this challenge, the dataset was split according to the identified broad cell types. Subsequently, each cell subset was individually filtered to exclude transcripts that occurred in fewer than 5 cells. *hg38 refseq-r80 10kb_up_and_down_tss.mc9nr.feather* and *hg38 refseq-r80 500bp_up_and_100bp_down_tss.mc9nr.feather* databases, as well as the human v9 motif collection were obtained from cisTarget (https://resources.aertslab.org/cistarget/). The GRNBoost2 algorithm was implemented for gene regulatory network inference, followed by the computation of enriched motifs and regulon predictions using the CLI ctx function. Finally, to estimate regulon activity scores in each cell, the aucell function was utilised. In the analysis of tumour cells, the workflow had one modification: SCENIC GRN inference was concurrently conducted for both tumour and PT PRAP1+ epithelial cells. This modification facilitated a direct comparison of TF activities between these two cell populations. Regulons demonstrating an absolute log2FC greater than 0.5 and an adjusted p-value below 0.05 were classified into activated and inactivated TF signatures.

### Data clustering and annotation

For each broad cell type, a total of 20 independent pySCENIC runs were performed. Results were subsequently filtered so that only regulons detected in a minimum of 19 out of 20 runs were retained. Meanwhile, the targets of the remaining TFs were filtered by selecting those that recurred in 19/19 or 20/20 runs. For each broad cell type subset, the resulting 20 filtered AUCell matrices were averaged to produce a single AUCell matrix, which was subsequently used to create a Seurat [67] object. Batch correction was performed using Seurat’s integration pipeline. Integration anchors were determined using the *FindIntegrationAnchors* function with default parameters and using all of the remaining regulons as features. An integrated Seurat object was obtained using the *IntegrateData* function with default parameters as well. Data was then scaled (*ScaleData*) and PCA performed (*RunPCA*), using all of the filtered regulons as features. Next, nearest-neighbour graph construction, cluster determination, and non-linear dimensionality reduction were carried out using *FindNeighbors*, *FindClusters* and *RunUMAP* functions, respectively. Differentially active regulons were identified using the *FindAllMarkers* function. The resulting cluster labels were then transferred to the original scRNA-seq Seurat object of the respective broad cell type, followed by scRNA-seq UMI count normalisation (*SCTransform*) and differential gene expression analysis (*FindAllMarkers*). Regarding tumour vs PT PRAP1+ differential expression analysis results, genes with an absolute log2FC value above 1 and an adjusted p-value below 0.05 were classified into up- and down-regulated signatures.

### Tumour meta-program determination and characterisation

The integrated Seurat [67] tumour cell object containing regulon activity scores (see above) was split according to the patient of origin. Next, for each patient, the corresponding AUCell matrix was scaled using the *ScaleData* function, followed by replacing all negative values in the matrix by zero. For each tumour, top 10 regulon activity modules were calculated using the *nmf* function of the NMF package (v0.26). For each regulon module, the top ten regulons with the highest weight were selected to define an intra-tumour activity program. The resulting regulon modules were then filtered to retain only those that had standard deviations above 0.2 among tumour cells. Finally, the resulting intra-tumour regulon activity modules were clustered based on pair-wise Jaccard index

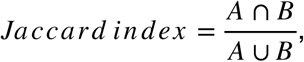

where A and B correspond to two intra-tumour regulon activity programs. This allowed the identification of regulon meta-programs that are shared across multiple tumours. Regulons shared by at least 50% of tumours with a particular meta-program were used to define that meta-program. Seurat *AddModuleScore* function with ‘nbin’ and ‘ctrl’ parameters being 6 and 50, respectively, was used to calculate the average activity levels of each meta-program in each tumour cell. Reactome [34] pathway enrichment analysis of the meta-programs was carried out using the standard workflow of the gprofiler2 [68] package.

### Collection of prognostic, ccRCC-associated, common essential, and drug target genes

Prognostic genes were retrieved from the Human Protein Atlas [36] (HPA) using the search term “Renal cancer”. The resulting list was further split into favourable and unfavourable prognosis gene signatures (FPGs and UPGs, respectively) based on survival analysis. ccRCC-associated genes (CAGs) were obtained from DisGeNET [35] using the following search terms: “Conventional (Clear Cell) Renal Cell Carcinoma”, “Clear-cell metastatic renal cell carcinoma”, and “Hereditary clear cell renal cell carcinoma”; only curated associations were considered. Common essential genes from the CRISPR and RNAi screens were sourced from the DepMap [37] database; the union of these two lists was used as a signature in downstream analyses. Clinical trial data was downloaded from the DrugBank [69] database, focusing on trials investigating potential treatments (instead of, for example, diagnostic, preventative, or supportive care). Furthermore, trials that were withdrawn, terminated, or suspended were excluded. Separately, only those target genes that have a ‘yes’ for DepMap’s pharmacological action section were included in the Class 1 set to ensure that the drug directly interacts with the target. For drug entries missing from the database or annotated as stubs, a literature search was conducted to identify their targets.

### Preprocessing and differential abundance testing of bulk omics data

Bulk ccRCC omics data from 110 donors were sourced from the Clark et al. [70] study. Across all omics layers, non-ccRCC samples (C3L-00359, C3N-00313, C3N-00435, C3N-00492, C3N-00832, C3N-01175, and C3N-01180) and their corresponding normal adjacent tissue samples, as identified by the original authors, were excluded from the analyses.

Bulk RNA-seq data were acquired using the TCGAbiolinks[71] package. During pre-processing, a contaminated transcriptomics sample (C3N-00314) was also excluded. Count data were filtered to retain transcripts with at least 10 counts in ≥50% of healthy or tumour samples. Library size normalization factors were then computed using the “TMM” method (*calcNormFactors*), and expression values were transformed with the *voom* function. For donors with multiple sequencing runs of the same tissue type (healthy or tumour), expression values were averaged. Differentially expressed genes were identified using the *lmFit* and *eBayes* functions of the limma [72] package. Genes with an absolute log2FC value above 2 and an adjusted p-value below 0.05 were used to define up- and down-regulated transcript signatures.

Copy number variation data were also obtained using the TCGAbiolinks[71] package. To identify genes with recurrent CNVs, we first excluded those located on the X and Y chromosomes. For donors with multiple sequencing runs, CNV values were averaged. Next, genes with missing values in more than 50% of samples were removed. Following guidelines from the Catalogue of Somatic Mutations in Cancer, (https://cancer.sanger.ac.uk/cosmic/help/cnv/overview), a gene was defined as amplified if present in ≥5 copies and as deleted if completely absent in a given donor. CNV frequencies were then calculated across all samples, and the top 1,000 genes most frequently affected by gains or losses were retained. This corresponded to a minimum of 7 donors for deletions and 26 donors for amplifications.

Proteomics and phosphoproteomics data were obtained from the CPTAC data portal (https://cptac-data-portal.georgetown.edu/cptac/s/S050; CPTAC3_CCRCC_Whole_abundance_gene_protNorm =2_CB.tsv; 6_CPTAC3_CCRCC_Phospho_abundance_phosphopeptide_protNorm=2_CB_imputed_12 11.tsv). For proteomics data, proteins with more than 50% of missing values were removed, while the remaining missing values were imputed using the DreamAI ensemble method (https://github.com/WangLab-MSSM/DreamAI). Differential protein and phosphopeptide abundance analyses were conducted utilising the *lmFit* and *eBayes* functions of the limma [72] package. Significant changes in protein levels in tumour tissues were identified with criteria of an absolute log2FC greater than 1 and an adjusted p-value below 0.05. For phosphoproteomics, a gene was considered differentially phosphorylated if at least one of its phosphosites had an absolute log2FC greater than 1 and an adjusted p-value below 0.05.

### Analysis of PPI network topology and CRISPR gene dependency

Human protein–protein interaction (PPI) data were sourced from the STRING [38] (v11.5) database and filtered to retain interactions with a combined confidence score of ≥400. Subsequently, the networkx [73] package was used to compute degree, betweenness, eigenvector, and PageRank centralities for each node within the network. Further, the networkx [73] package facilitated the calculation of shortest path lengths within the network. This analysis entailed computing both the average and minimum shortest path distances from a given node to every gene in a particular signature. In the context of the machine learning feature space, minimum distance to DTGs and ‘Other’ genes were excluded to prevent label leakage and avoid biasing predictions towards genes with a minimum distance to Others greater than 0, respectively. Lastly, for each gene, the PPI network was used to calculate the ratios of its PPI neighbours that belong to a particular signature.

CRISPR gene dependency data were obtained from the DepMap [37] database, followed by filtering to retain all available (n=16) ccRCC cell lines. Missing dependency values for a given gene in a specific cell line were imputed using the mean dependency score across cell lines with non-missing values for that gene. Average gene dependency scores of all PPI neighbours of a particular gene were calculated for each cell line individually.

### Machine learning methodology: framework, model optimisation, and performance evaluation

To estimate the therapeutic potential of individual genes, we applied conventional machine learning methods for binary classification. The 199 known drug target genes (DTGs) were assigned as the positive class (Class 1) and split into a training and cross-validation set (80%, n = 159) and an independent test set (20%, n = 40) held out for final model evaluation. The negative class (Class 0) comprised the remaining gene pool (n = 17,681), which was similarly divided into training (80%, n = 14,151) and test (80%, n = 3,530) sets. Here, the assumption was that most genes fulfil roles other than as therapeutic targets, thereby minimising the incidence of false negatives within the training data. To create balanced training sets and reduce model variance through averaging, the 14,151 negative training samples were partitioned into 89 non-overlapping subsets of 159 genes each, matching the number of positive training samples. Therefore, each balanced training set consisted of the same 159 positive samples and a unique set of 159 negative samples, ensuring every negative gene was sampled exactly once across all training iterations.

Subsequently, we applied five machine learning classifiers: Logistic Regression (LR), Support Vector Machine (SVM), Random Forest (RF), Gradient Boosting (GB), and an ensemble model that averaged the predicted Class 1 probabilities across LR, SVM, RF, and GB methods. For SVM, hyperparameter tuning was conducted using grid search with 5-fold cross-validation, optimising the kernel function (linear or RBF), cost parameter, and kernel bandwidth (for the RBF kernel). For RF, an initial grid search determined the optimal forest size, followed by a randomised search to fine-tune the number of features considered for node splitting, maximum tree depth, and minimum sample thresholds for node splits and leaf nodes. Similarly, the GB model’s learning rate and forest size were optimised via grid search, with a subsequent randomised search for decision tree parameters (analogous to RF). All five models were trained on each of the 89 balanced training sets. Performance was assessed using 5-fold cross-validation based on standard classification metrics: accuracy, precision, recall, F1 score, and area under the receiver operating characteristic curve (AUROC).

### Model testing on unseen data

Upon establishing the ensemble classifier as the best-performing method based on cross-validation, we proceeded to evaluate its performance on unseen data. Mirroring the approach employed for training procedure, we generated 88 balanced test sets. This involved partitioning the 3,530 negative class genes reserved for testing into groups of 40 —matching the number of held-out positive samples. Each negative group was paired with the same set of 40 positive test samples to form the test sets. Each of the 89 ensemble models was then evaluated on all 88 test sets using accuracy, precision, recall, F1 score, and AUROC metrics. Final performance was calculated by averaging results across all ensemble models and test sets.

### Spatial transcriptomics data analysis

Tumour core and tumour-normal interface spatial transcriptomics data was obtained from Mendeley Data (https://data.mendeley.com/datasets/g67bkbnhhg/1). The Cell2location[74] package was implemented to map cell clusters from scRNA-seq data to the 10X Genomics Visium spatial transcriptomics measurements. First, basic filtering of spots was carried out using Scanpy’s[75] *pp.filter cells* function to remove spots with transcript counts below 2000 or above 35,000, less than 500 detected genes, and a mitochondrial gene fraction exceeding 20%. Additionally, slides 6800STDY12499504 and 6800STDY12499505 were also removed due to having fewer than 500 spots. Next, scRNA-seq data was filtered using Cell2location’s *filter_genes* function with the following parameters: ‘cell_count_cutoff’ = 5, ‘cell_percentage_cutoff2’ = 0.01, ‘nonz_mean_cutoff’ = 1.05. The unnormalised mRNA count matrix was used as input for this filtering step, after which 16285 genes and 250331 cells remained in the dataset. Next, a negative binomial regression model was implemented to estimate the reference signature of cell types detected in the scRNA-seq dataset. Here, patient IDs were used as batch keys, and the model was trained using the *mod.train* function with the following parameters: ‘max_epochs’ = 100, ‘batch_size’ = 2500, ‘train_size’ = 1, ‘lr’ = 0.002. The summary of the posterior distribution was exported using the *mod.export_posterior* function. The resulting reference signature model was then used to predict the spatial abundance of cell types. Only genes identified in both scRNA-seq and spatial transcriptomics were retained. Cell type abundance was estimated separately for the tumour core and tumour-normal interface slides, employing the following parameters: ‘N_cells_per_location’ = 20, ‘detection_alpha’ = 200, ‘max_epochs’ = 30,000.

### In vitro experiments

#### Cell lines

A498, 769-P and 786-O cell lines were kindly provided by the Sakari Vanharanta laboratory. Cells were expanded using R10 medium (RPMI1640 (gibco), 10% Fetal Bovine Serum (Sigma) 1% Penicillin-Streptomycin (Sigma) and 2mM L-Glutamine (gibco). Cells were expanded using adherent T75s in 37°C, 5% CO2 incubators with Accutase (Invitrogen) passaging. All cells used for experiments were below passage 15.

#### Drugs

ML385 (SML1833-5MG), BLT-1 (373210-25MG), POMHEX (HY-131904-5MG), ERW1041E (5095220001), LRRK2-IN-1 (Biotechne Tocris, #4273), and HMOX1-IN-1 (TA9H94532D16-5MG) powders were solubilised in DMSO (Invitrogen). BAPN (S5340-25MG) was solubilised in ddH2O. All reconstituted solutions were stored as directed by the manufacturer.

#### Cytotoxicity

5000 proliferating cells in 25µL of R10 medium was added to wells of pre-warmed adherent 384 well plates. After 24 hours at 37°C, 5% CO2, R10 medium was aspirated before 25µL drug-supplemented R10 medium or DMSO controls were added. Supplemented-R10 was incubated with cells for 2 days, before 25µL CellTiterGlo3D Viability Assay (ProMega) administration and viability determination through luminescence recording by FLUOstar Optima (BMG Labtech). Drugs were assessed across two experimental rounds: round 1 examined ML385, BLT-1, POMHEX, ERW1041E, and BAPN, while round 2 examined LRRK2-IN-1 and HMOX1-IN-1.

#### Proliferation

4000 or 2500 proliferating cells (A498/769P or 786O) were seeded per 96 well plate well in 100µL of drug or control supplemented R10 medium. Plates were equilibrated for 2hrs at 37°C, 5% CO2, before 10µL CCK8 (abcam) was added with incubation for 4hr, 37C, 5% CO2 and the recording of absorbance at 450nm using the FLUOstar Optima (BMG Labtech). CCK8 testing was repeated on separate plates on days 1, 2 and 3, with drug supplemented medium-only controls included for each timepoint to account for background CCK8 reduction. Drugs were assessed across two experimental rounds: round 1 examined ML385, BLT-1, POMHEX, ERW1041E, and BAPN, while round 2 examined LRRK2-IN-1 and HMOX1-IN-1. For plotting purposes, to account for minor differences in baseline vehicle proliferation between experimental rounds, OD450 values from round 2 were rescaled such that the mean background-corrected vehicle OD450 per cell line and time point matched the corresponding values from round 1.

#### Statistical analyses

All data are presented as the mean ± standard error of the mean (SEM) or standard deviation (SD). Statistical analyses and IC50 value calculations were performed in GraphPad Prism 10.2.3. For comparisons involving more than three groups, statistical significance was determined using the Kruskal-Wallis test followed by Dunn’s multiple comparisons test.

