## Supplementary Figures for "A network medicine framework for multi-modal data integration in therapeutic target discovery"

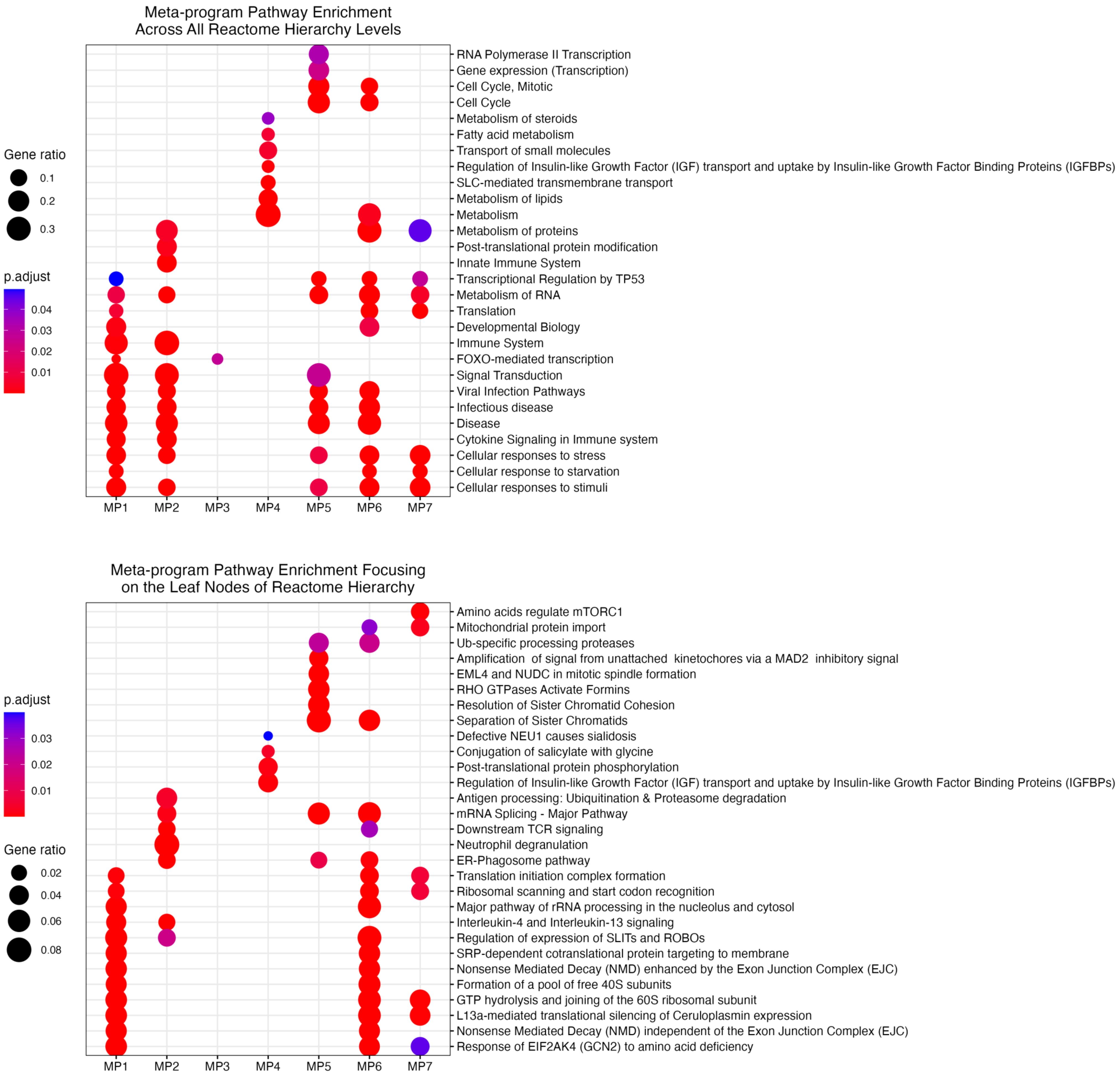


**Supplementary Figure 1: Reactome pathway enrichment analysis of tumour meta-programs.** Transcription factors and their target genes defining each meta-program were used to perform Reactome pathway enrichment analyses. Top panel: analysis results incorporating pathways at all levels of the Reactome pathway hierarchy. Bottom panel: enrichment analysis focused exclusively on the terminal (leaf) nodes of the hierarchy, ensuring specificity of functional annotations.

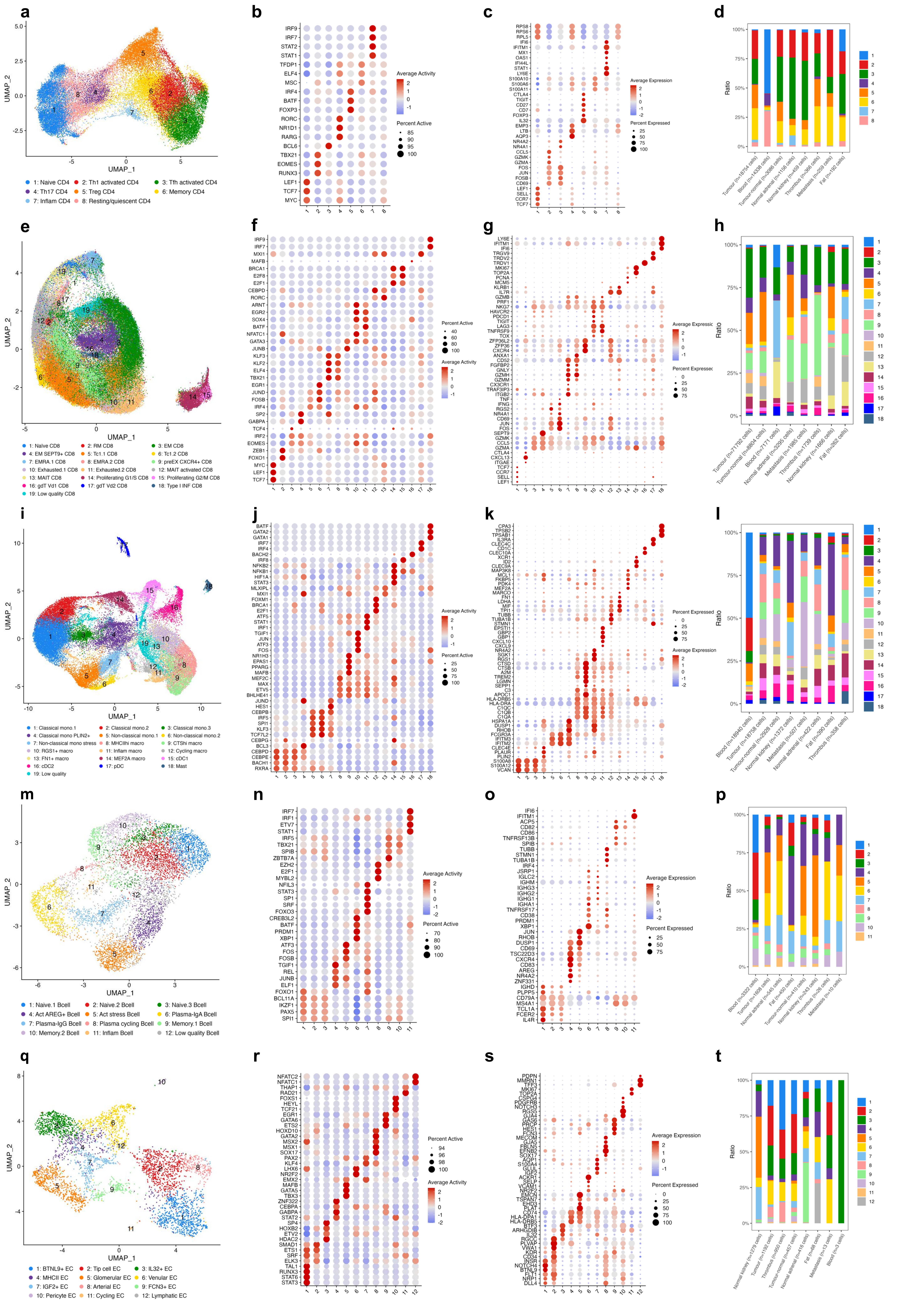

a
b
c
d
UMAP_1
UMAP_2
1: Naive CD4
2: Th1 activated CD4
3: Tfh activated CD4
4: Th17 CD4
5: Treg CD4
6: Memory CD4
7: Inflam CD4
8: Resting/quiescent CD4
Average Activity
Percent Active
Average Expression
Percent Expressed
Ratio
Tumour (n=18754 cells)
Blood (n=14338 cells)
Tumour-normal (n=3086 cells)
Normal adrenal (n=1156 cells)
Normal kidney (n=459 cells)
Thrombus (n=366 cells)
Metastasis (n=259 cells)
Fat (n=192 cells)
e
f
g
h
1: Naïve CD8
2: RM CD8
3: EM CD8
4: EM SEPT9+ CD8
5: Tc1.1 CD8
6: Tc1.2 CD8
7: EMRA.1 CD8
8: EMRA.2 CD8
9: preEX CXCR4+ CD8
10: Exhausted.1 CD8
11: Exhausted.2 CD8
12: MAIT activated CD8
13: MAIT CD8
14: Proliferating G1/S CD8
15: Proliferating G2/M CD8
16: gdT Vd1 CD8
17: gdT Vd2 CD8
18: Type I INF CD8
19: Low quality CD8
Tumour (n=71792 cells)
Tumour-normal (n=8804 cells)
Blood (n=7171 cells)
Normal adrenal (n=3295 cells)
Metastasis (n=1985 cells)
Thrombus (n=1739 cells)
Normal kidney (n=1666 cells)
Fat (n=262 cells)
i
j
k
l
1: Classical mono.1
2: Classical mono.2
3: Classical mono.3
4: Classical mono PLIN2+
5: Non-classical mono.1
6: Non-classical mono.2
7: Non-classical mono stress
8: MHCIIhi macro
9: CTShi macro
10: RGS1+ macro
11: Inflam macro
12: Cycling macro
13: FN1+ macro
14: MEF2A macro
15: cDC1
16: cDC2
17: pDC
18: Mast
19: Low quality
Blood (n=18940 cells)
Tumour (n=18758 cells)
Tumour-normal (n=2928 cells)
Normal kidney (n=1372 cells)
Metastasis (n=507 cells)
Normal adrenal (n=422 cells)
Fat (n=390 cells)
Thrombus (n=358 cells)
m
n
o
p
1: Naive.1 Bcell
2: Naive.2 Bcell
3: Naive.3 Bcell
4: Act AREG+ Bcell
5: Act stress Bcell
6: Plasma-IgA Bcell
7: Plasma-IgG Bcell
8: Plasma cycling Bcell
9: Memory.1 Bcell
10: Memory.2 Bcell
11: Inflam Bcell
12: Low quality Bcell
Blood (n=3322 cells)
Tumour (n=1608 cells)
Normal adrenal (n=545 cells)
Fat (n=450 cells)
Tumour-normal (n=410 cells)
Normal kidney (n=243 cells)
Thrombus (n=26 cells)
Metastasis (n=10 cells)
q
r
s
t
1: BTNL9+ EC
2: Tip cell EC
3: IL32+ EC
4: MHCII EC
5: Glomerular EC
6: Venular EC
7: IGF2+ EC
8: Arterial EC
9: FCN3+ EC
10: Pericyte EC
11: Cycling EC
12: Lymphatic EC
Normal kidney (n=1279 cells)
Tumour (n=1192 cells)
Thrombus (n=950 cells)
Tumour-normal (n=457 cells)
Normal adrenal (n=416 cells)
Fat (n=68 cells)
Metastasis (n=13 cells)
Blood (n=3 cells)

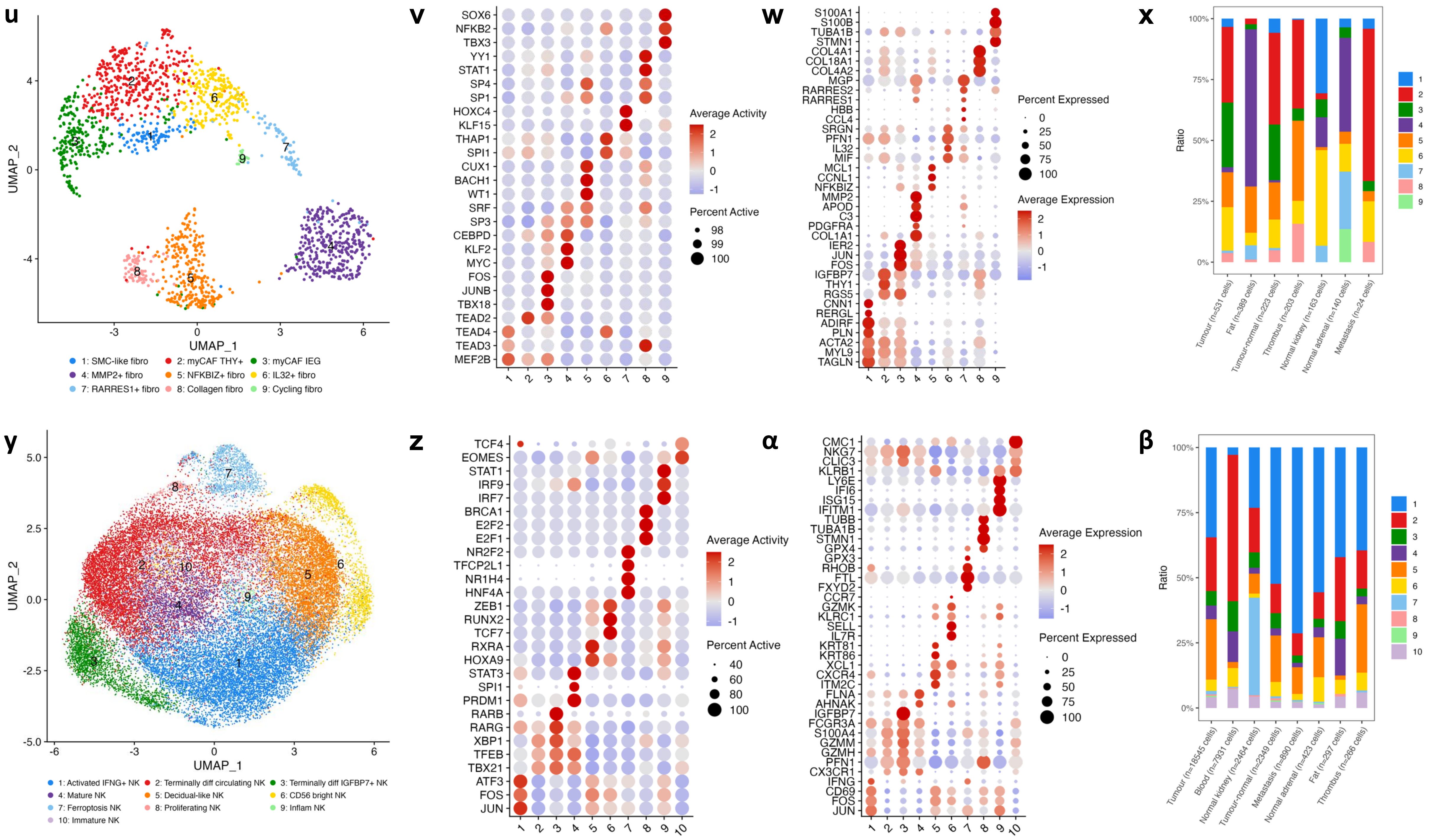


**Supplementary Figure 2: Transcription factor activity-based sub-clustering of CD4, CD8, myeloid, B, epithelial, fibroblast, and NK cell compartments.**
**a, e, I, m, q, u, y** UMAPs depicting CD4, CD8, myeloid, B, epithelial, fibroblast, and NK cell sub-clustering. **b, f, j, n, r, v, z** Regulon (TF activity) scores across the identified sub-populations. **c, g, k, o, s, w, α** Canonical marker gene expression plots. **d, h, l, p, t, x, β** Sub-cluster distributions across anatomical regions sampled for sequencing.

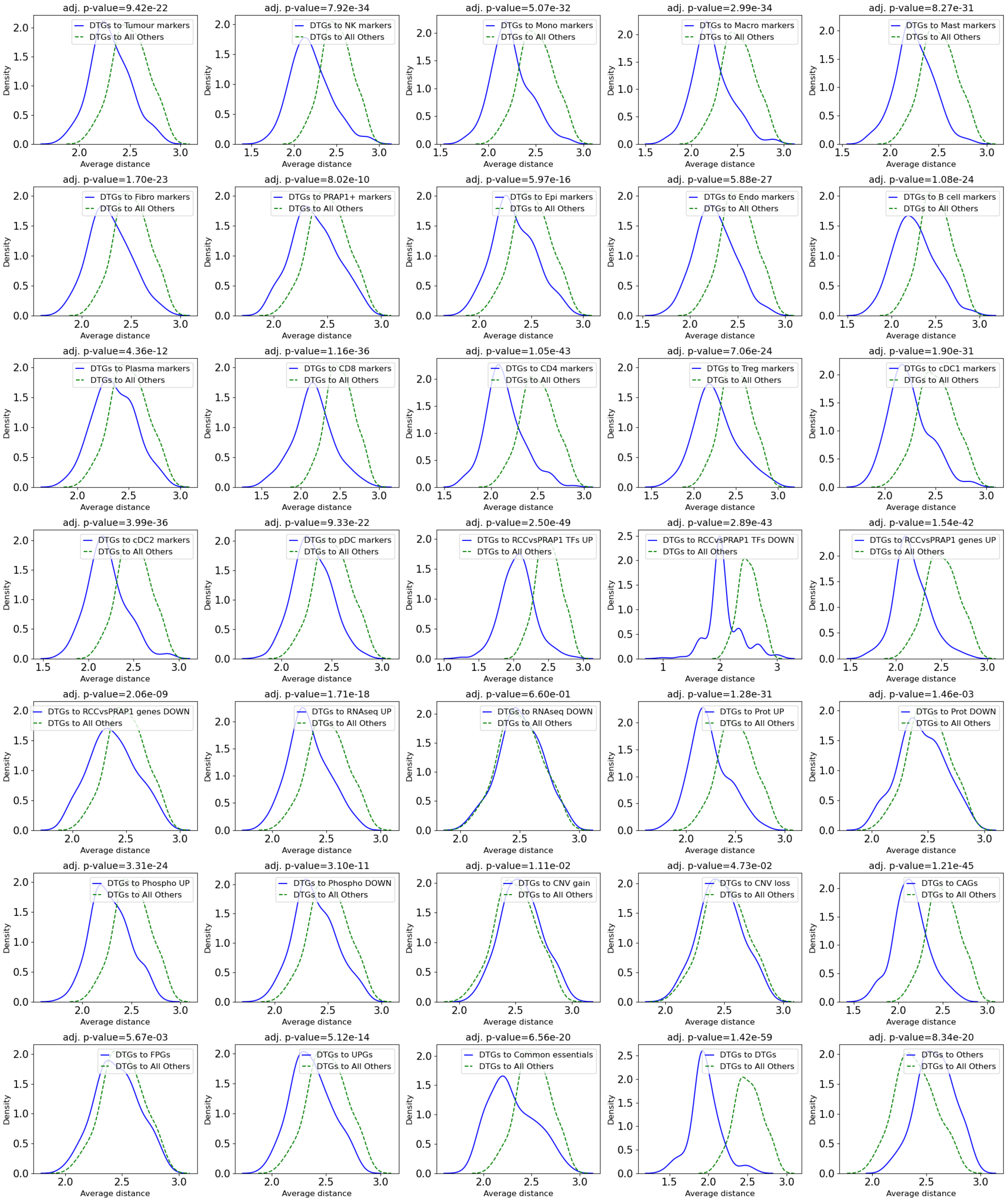


**Supplementary Figure 3: Proximity of drug target genes to curated signatures within the PPI network.** Average shortest path distance distributions within the PPI network are compared to assess the proximity of drug target genes (DTGs) to each curated gene signature (blue) relative to all remaining network nodes (green). Statistical significance was assessed using BH-adjusted Wilcoxon rank-sum tests; adjusted *p*-values are indicated.

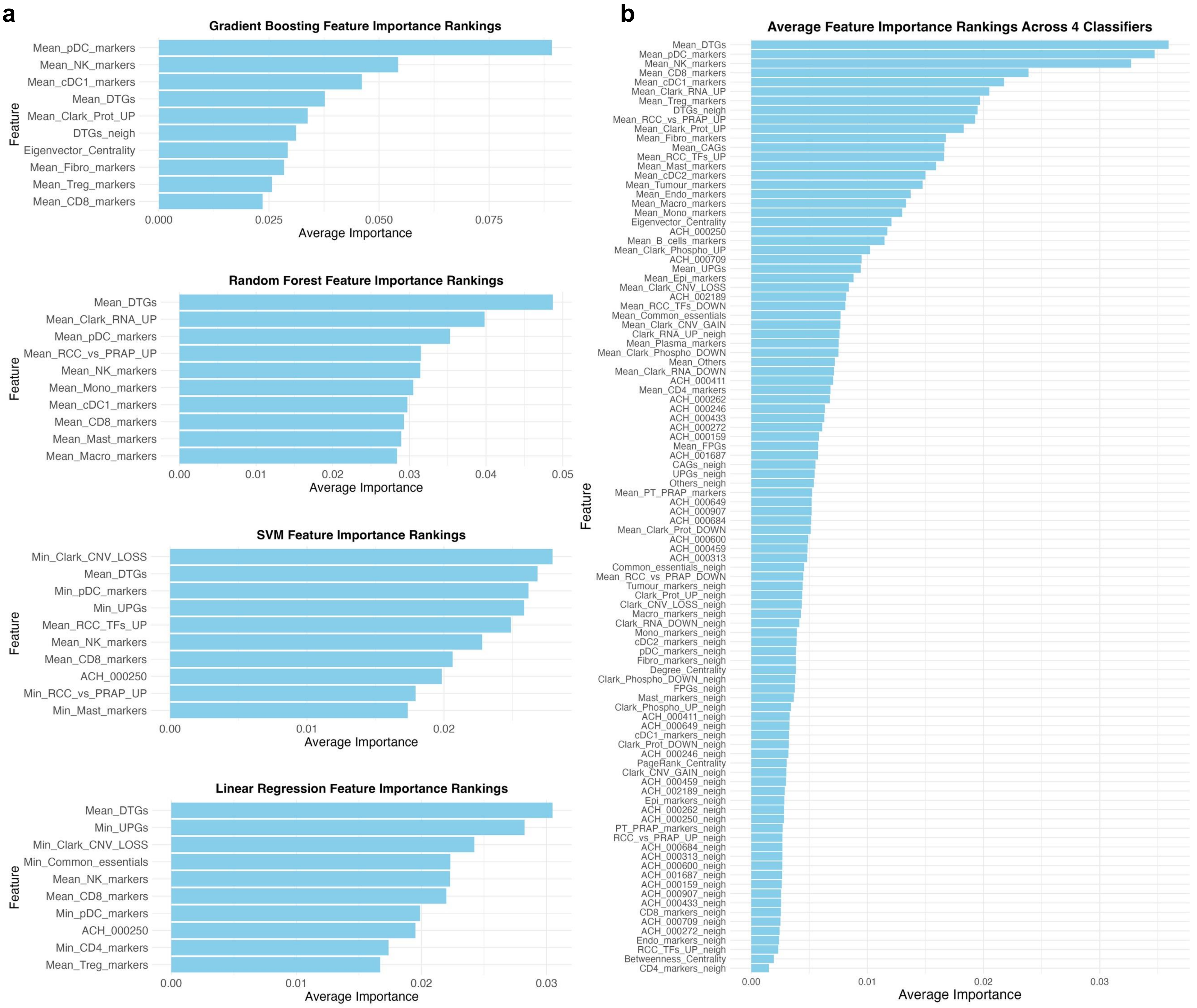


**Supplementary Figure 4: Feature importance ranking across machine learning classifiers.**
**a** Top 10 most important features per classifier. Feature importance values were normalised to sum to 1, such that the *x*-axis reflects the relative contribution of each feature to model predictions. **b** Top 100 most informative features, ranked by average importance scores across four machine learning classifiers. Feature importance scores were similarly normalised to represent relative importance, with the *x*-axis indicating the proportion of total importance attributed to each feature.

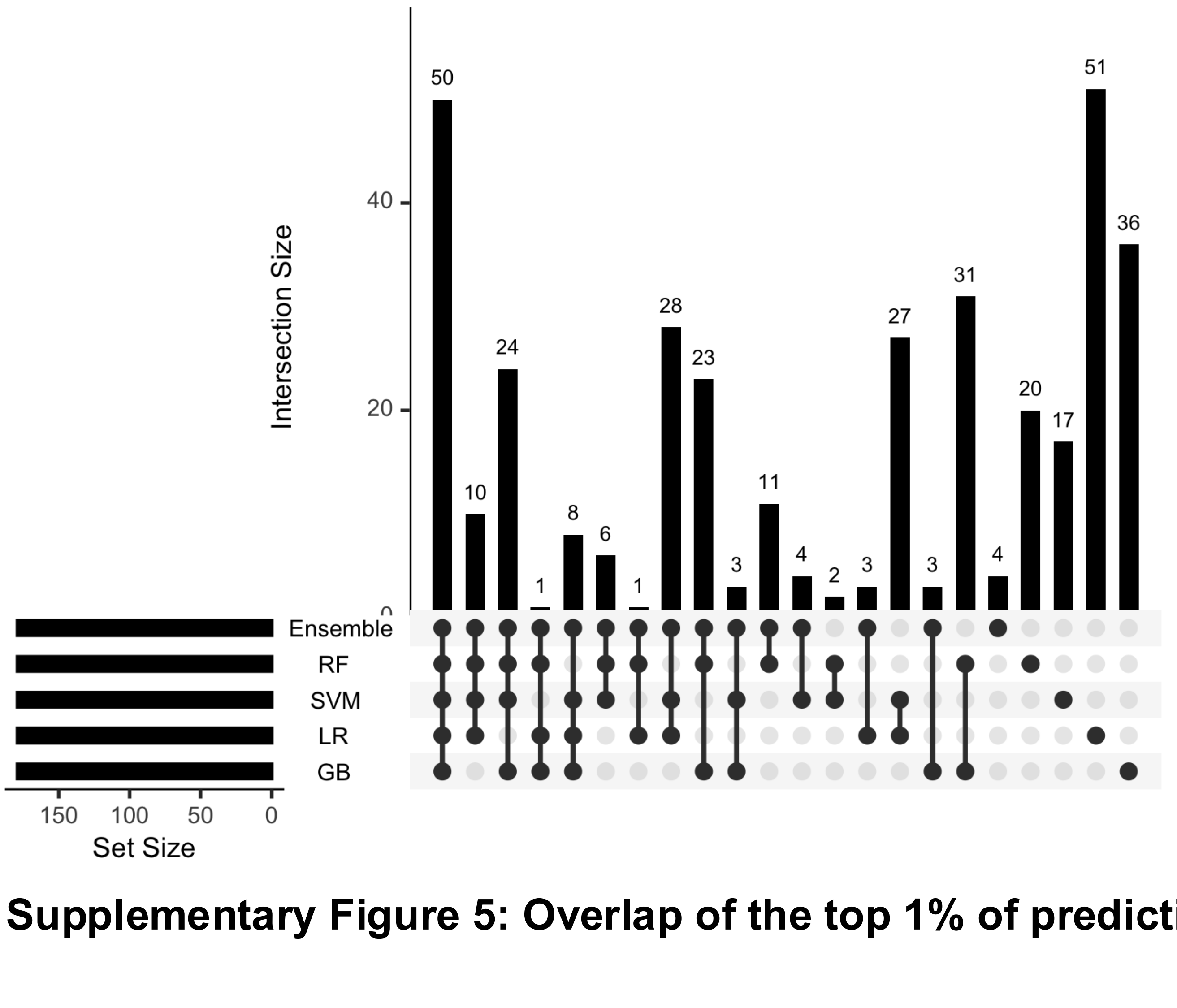


**Supplementary Figure 5: Overlap of the top 1% of predictions across ML classifiers.**

a

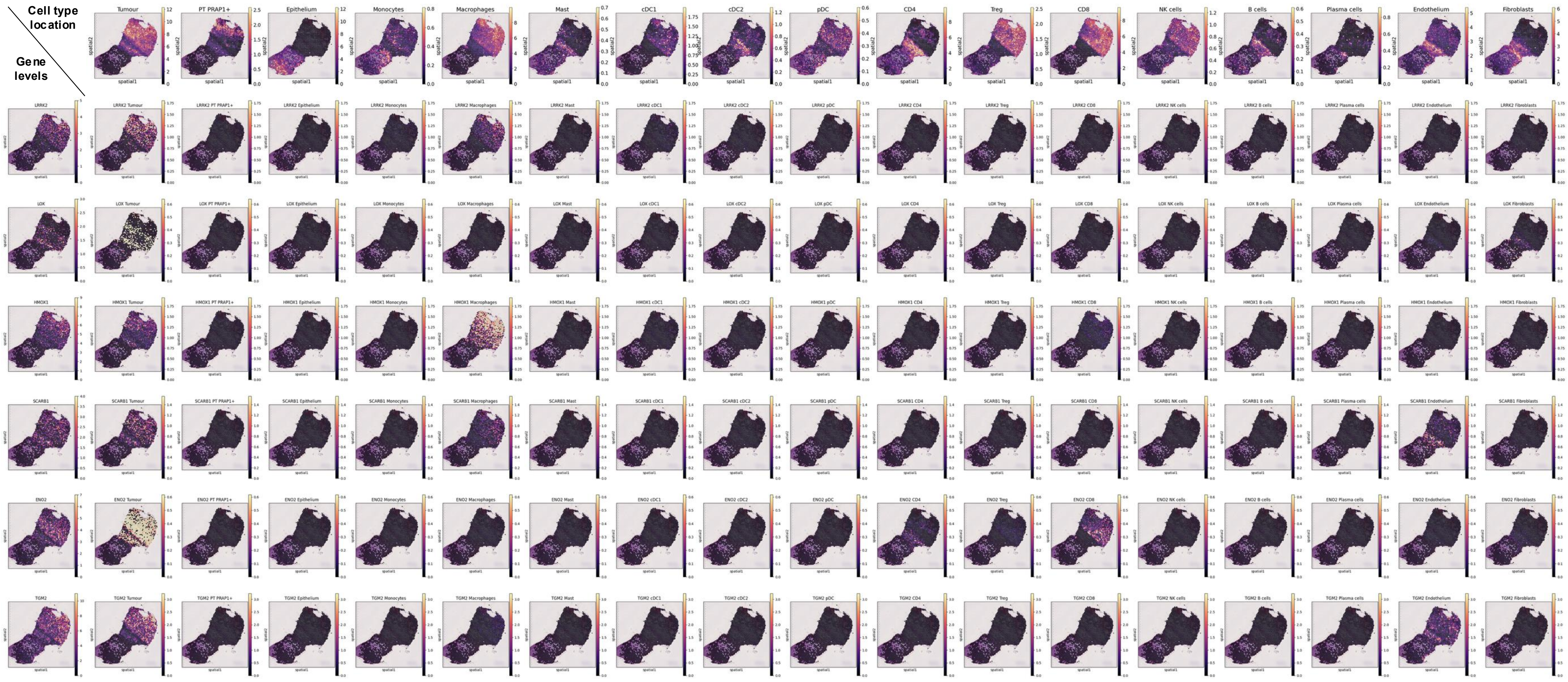


b

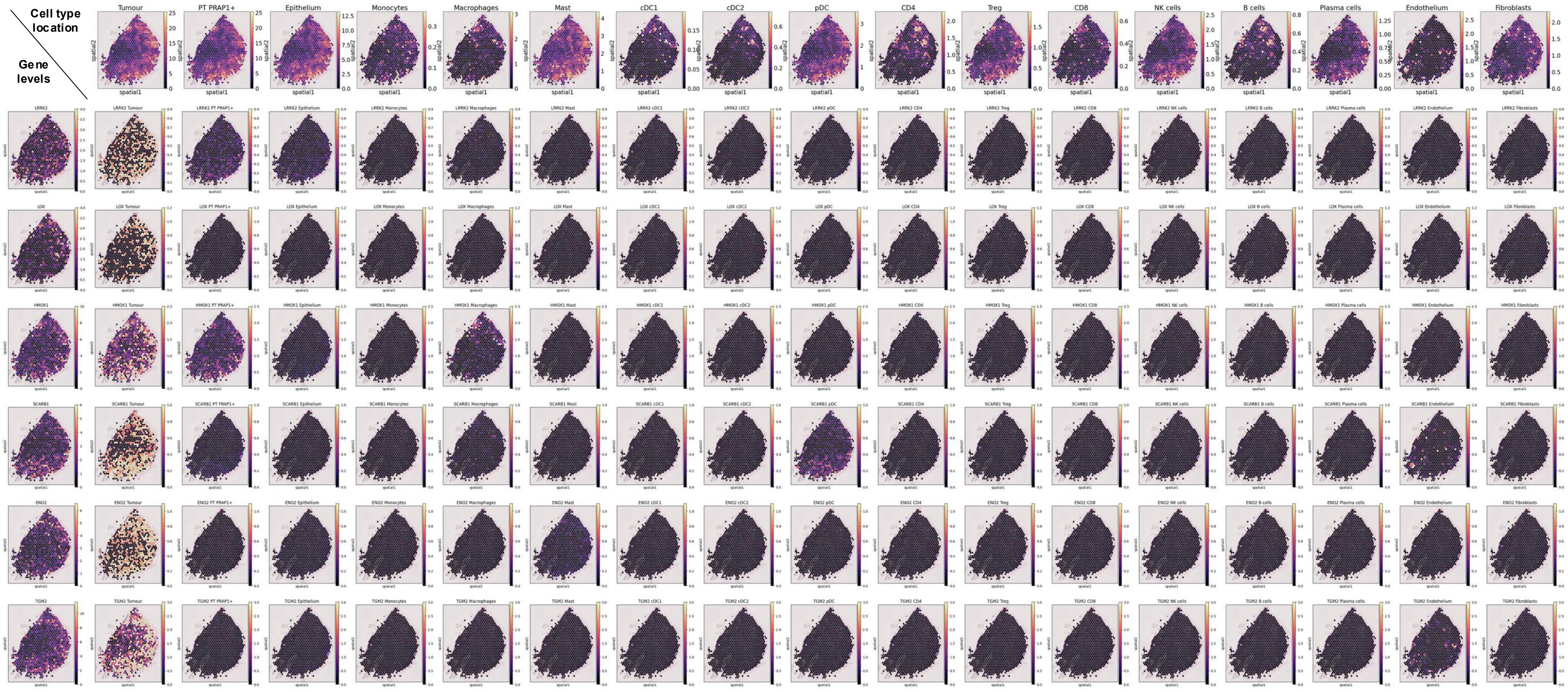


**Supplementary Figure 6: Target expression patterns as observed in ccRCC tumour spatial transcriptomics data.**

**a** ccRCC tumour-normal interface spatial transcriptomics data from patient PD47171. The top row depicts localisation of a given cell type, while the subsequent rows show gene levels across cell types. **b** ccRCC tumour core spatial transcriptomics data from patient PD43948. The top row depicts localisation of a given cell type, while the subsequent rows show gene levels across cell types.

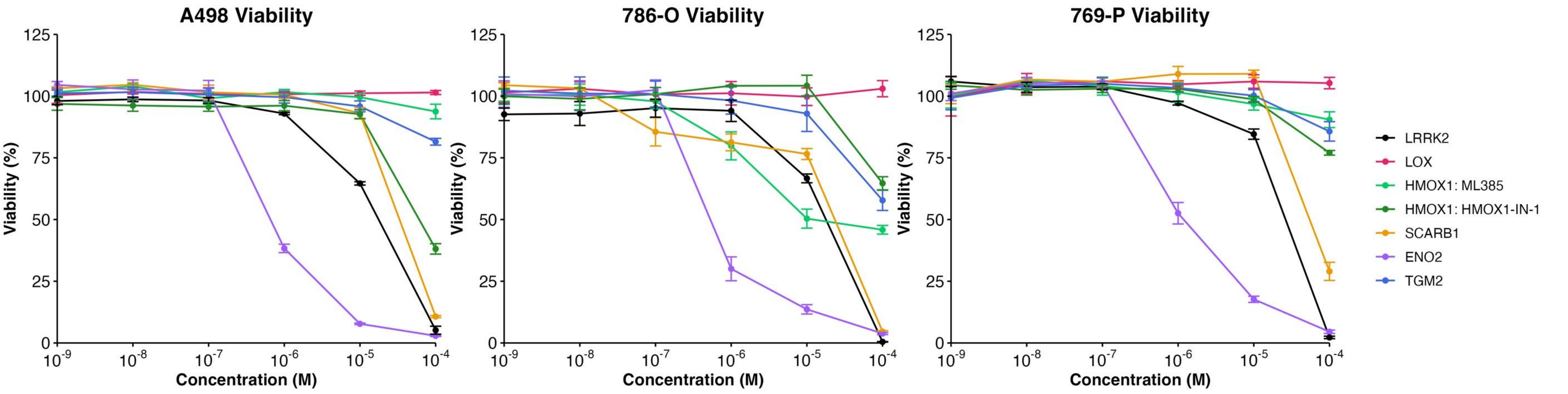


**Supplementary Figure 7: Inhibitor cytotoxicity across ccRCC cell lines.** A498, 786-O, and 769-P cells were treated in triplicate for 48 hours with either vehicle controls or small molecule inhibitors across six concentrations. Results are presented as mean± SD of the three technical replicates.

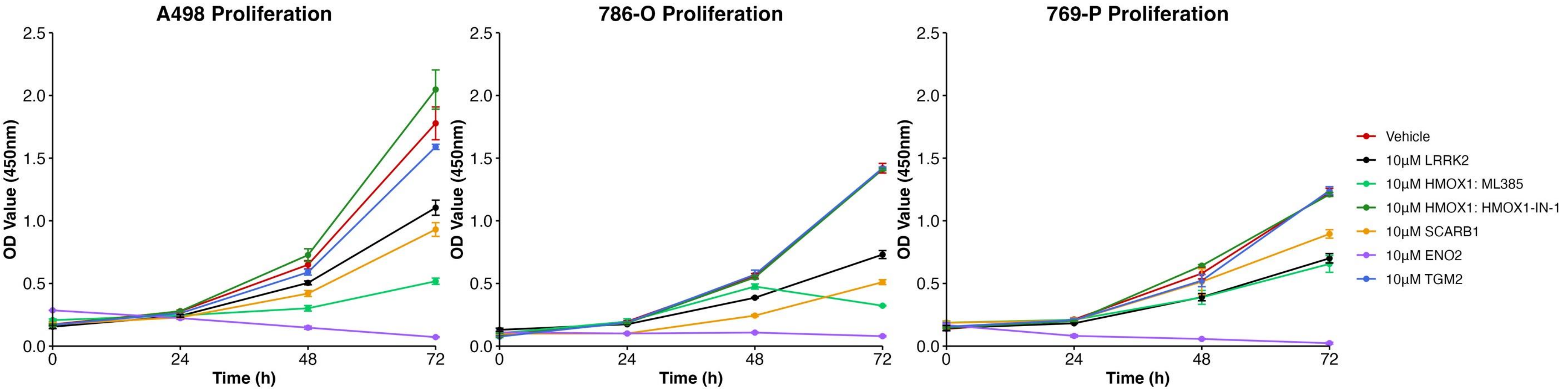


**Supplementary Figure 8: Inhibitor cytostatic effects across ccRCC cell lines.** A498, 786-O, and 769-P cells were treated in triplicate for 72 hours with either vehicle controls or 10µM of inhibitors targeting the genes of interest. Results are presented as mean ± SD of the three technical replicates.

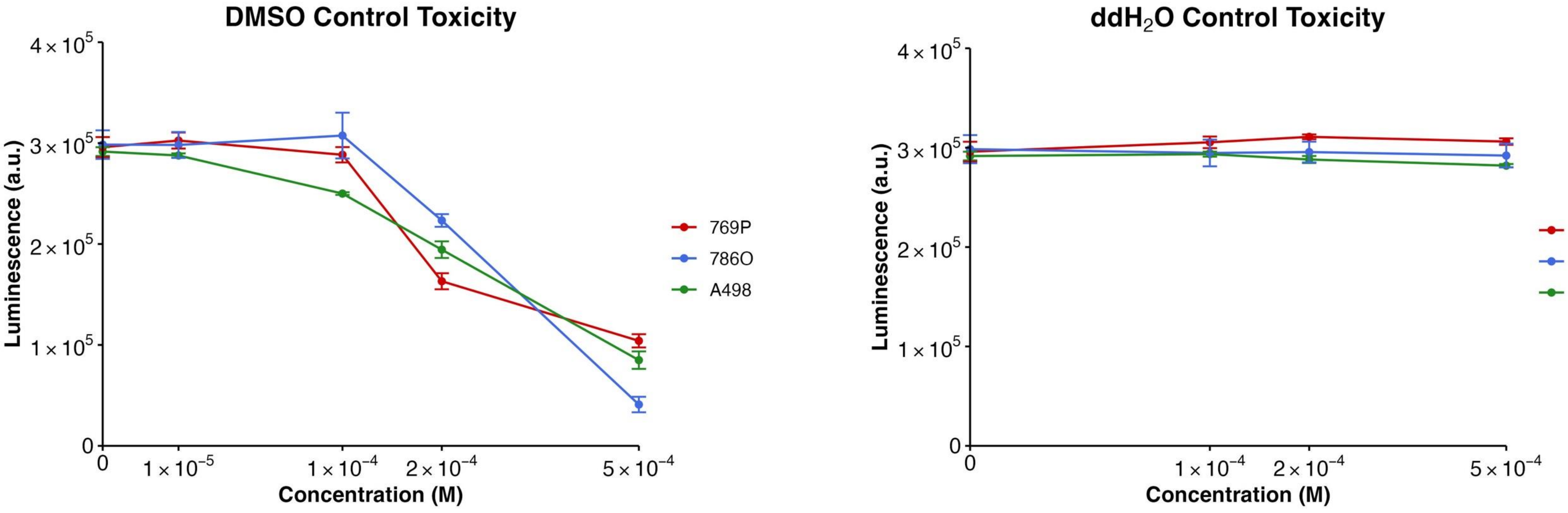


**Supplementary Figure 9: DMSO and ddH2O carrier cytotoxicity across ccRCC cell lines.**
